# Filament assembly, stabilisation, and ATP-dependent RAD51 filament capping and remodeling by RAD51C-XRCC3 complex

**DOI:** 10.64898/2026.11.20.768045

**Authors:** Ondrej Belan, Mario Spirek, Gareth J. Williams, Michaela Bednarova Pospisilova, Jarmila Mlcouskova, Lucia Molnarova, Jordan Pepper, Simon J. Boulton, John A. Tainer, Zbynek Prokop, Lumir Krejci

## Abstract

Homologous recombination requires tightly controlled assembly and maturation of RAD51 nucleoprotein filament, yet the mechanistic roles of the RAD51 paralogs remain poorly understood. Here, we show that the human RAD51 paralog complex RAD51C-XRCC3 (CX3) remodels oligomeric RAD51 and promotes filament assembly while capping the 5’ filament end to stabilise it. CX3 protects RAD51 filaments from dissociation induced by competitor DNA or BRCA2-derived BRC repeats and converts them into a nuclease-sensitive remodelled state. This transition requires ATP-bound RAD51 and ATP hydrolysis by RAD51C, without altering RAD51 ATPase activity. ATP-binding-defective RAD51 fail to undergo remodeling and instead trigger a futile CX3 ATP turnover, suggesting coupling between RAD51 conformation and CX3 activity. Together, our findings establish CX3 as an ATP-dependent RAD51 filament assembly and remodelling factor and define a mechanism of filament maturation distinct from BRCA2- and BCDX2-mediated RAD51 regulation.

## INTRODUCTION

Homologous recombination (HR) is an essential pathway for the repair of DNA double-strand breaks (DSBs) and the maintenance of genome stability. In addition to its role in DSB repair, HR is essential for the protection, reversal and restart of stalled replication forks^1–3^. In response to replication stress, stalled forks are frequently reversed into a chicken foot-like structure promoted and stabilised by RAD51 nucleoprotein filament^2–4^. These reversed forks are bound by RAD51, thereby protecting the nascent strand from nucleolytic degradation by MRE11 and other nucleases^2,3,5–7^. The importance of stable RAD51 filament formation in protecting reversed forks is highlighted by the inability of disease-associated RAD51 mutants to prevent fork degradation in response to replication stress^6,8^. Furthermore, broken replication forks are rescued via a RAD51/BRCA2-mediated HR mechanism, which serves as an important barrier to prevent carcinogenesis^2^.

Central to RAD51 function is its ability to assemble into a helical nucleoprotein filament on ssDNA, a process tightly regulated by recombination mediator proteins. BRCA2, a tumour suppressor frequently mutated in hereditary breast and ovarian cancer, is the most extensively characterised HR mediator in human cells. BRCA2 interacts with RAD51 primarily through eight conserved BRC repeats, directly contacting ATPase domain of RAD51 and sequestering it in a monomeric state^9^. Beyond recruitment, BRCA2 facilitates RAD51 nucleation on RPA-coated ssDNA and stabilises the RAD51 filament by inhibiting RAD51’s intrinsic ATPase activity, thereby reducing filament turnover^10^. Cells lacking BRCA2 display hypersensitivity to DNA damaging agents and increased replication stress, along with impaired RAD51 focus formation and genomic instability^11^.

The five RAD51 paralogs, including RAD51B, RAD51C, RAD51D, XRCC2, and XRCC3, share structural homology with RAD51 but lack strand-exchange activity^12–14^. They organise into two principal complexes: the BCDX2 complex (RAD51B-RAD51C-RAD51D-XRCC2)^15,16^, which was initially suggested to act at early stage of presynaptic filament assembly by facilitating RAD51 nucleation on RPA-coated ssDNA^17^, and the CX3 complex (RAD51C-XRCC3)^15,16^ that was suggested to act downstream of RAD51 nucleation^17^. Analogously, during DNA replication, BCDX2 complex was proposed to act in upstream fork reversal and fork protection, while CX3 complex was proposed to play a role downstream in HR-mediated restart of stalled forks^18^. Recent structural and biochemical studies have revised this model. Structural analyses have provided mechanistic insight into BCDX2 architecture and revealed its role in directly promoting RAD51 filament nucleation and extension on RPA-coated ssDNA, as well as replication fork protection^19–21^. Our laboratory has further shown that BCDX2 efficiently promotes RAD51 nucleation on short ssDNA substrates, conditions relevant to processed DSB ends and stalled replication forks where ssDNA length is limiting^22^. Previous work on CX3 established conditions for purifying non-aggregated human CX3 and determined the crystal structure of the *Alvinella pompejana* CX3 core complex^23^. Structure-guided analysis of cancer-associated RAD51C variants, combined with DNA-fibre assays, revealed distinct requirements for replication fork protection and restart, including variants that retained homologous recombination proficiency despite exhibiting replication-associated defects^23^.

More recent work has suggested that the CX3 heterodimer can associate with DX2 (RAD51D-XRCC2) and form a functional heterotetrameric complex (X3CDX2) that stably caps the 5′-termini of assembled RAD51 filaments and promotes downstream strand invasion^12–14^. Lastly, the most recent structural cryo-EM analysis demonstrated that all five paralogs can co-assemble into a BCDX2-CX3 supercomplex coordinating filament assembly and stabilisation^13^. Additional cryo-EM studies demonstrated that RAD51 paralog exchange dynamically remodels these assemblies into filament-compatible intermediates that facilitate RAD51 loading on RPA-coated ssDNA^14^. Deficiency in any individual paralog results in impaired RAD51 focus formation, hypersensitivity to DNA interstrand crosslinks, and genomic instability^24,25^. The clinical importance of these factors is highlighted by biallelic mutations in BRCA2 and RAD51C causing Fanconi anaemia^26–28^, while monoallelic mutations in RAD51C, RAD51D, and RAD51B are associated with predisposition to breast and ovarian cancers^29–32^.

Parallels can be drawn from studies in *Caenorhabditis elegans*, where the RAD-51 paralog complex RFS-1/RIP-1 binds to the 5′ end of presynaptic RAD-51 filaments and remodels it to a more stable, flexible conformation, facilitating strand exchange^33,34^. These studies have raised several key questions including whether these activities are conserved outside of nematodes, what is the role of ATP binding and/or hydrolysis during this transition, and which of the human paralog complexes possess the filament remodeling activity. Furthermore, while the function of BCDX2 and X3CDX2 has been extensively characterized, whether CX3 can modulate RAD51 filaments assembly and/or stability is currently unclear.

Here, we provide detailed mechanistic analysis of human RAD51 filament regulation mediated by the CX3 complex. We show that CX3 binds RAD51 with high affinity, inducing rearrangements of its oligomeric state and promoting filament assembly and stability. CX3 binds 5’ filament end, consistent with a capping function, remodels it to a more open and stable conformation that is resistant to dissociation by the BRC repeat of BRCA2. These activities require ATP-bound RAD51 and ATP hydrolysis by CX3. In contrast, a RAD51 mutant deficient in ATP binding still allows capping by CX3, but this interaction fails to induce conformational change at the filament end and instead triggers a futile cycle of ATP hydrolysis by CX3. Together, these results provide novel and unexpected mechanistic insight into how the CX3 complex binds and promotes formation of RAD51 filament with altered and more stable conformation at the 5’ filament end, which may underlie its role in HR and genome maintenance.

## RESULTS

### CX3 directly binds RAD51 and associates with RAD51 nucleoprotein filament

We expressed and purified the human CX3 complex as a maltose binding protein (MBP) fusion from insect cells (**Fig. S1A**) using our optimised purification protocol combining a solubilising fusion tag with inclusions of nucleotide cofactor, which yields a native, soluble, and untagged human CX3 complex described in^23^.

Given previous yeast two-hybrid evidence for an interaction with RAD51^35^, we tested direct association using purified proteins. Pull-down assay demonstrated that CX3 strongly binds RAD51, irrespective of nucleotide cofactor present (**Fig. 1A and S1B**). Next, we monitored the mobility of RAD51-CX3 complex on native agarose gel. Using fluorescently labelled RAD51 we detected slow migrating protein species upon addition of CX3 to RAD51 (**Fig. S1C, D**). To quantify this interaction, we used biolayer interferometry (BLI), where His_6_-tagged CX3 was immobilised on a Ni-NTA-conjugated biosensor, and RAD51 association and dissociation were monitored in real time. RAD51 bound CX3 in a concentration dependent manner with apparent dissociation constant (*K*_D_) of 323 nM (**Fig. 1B**).

**Fig. 1:**
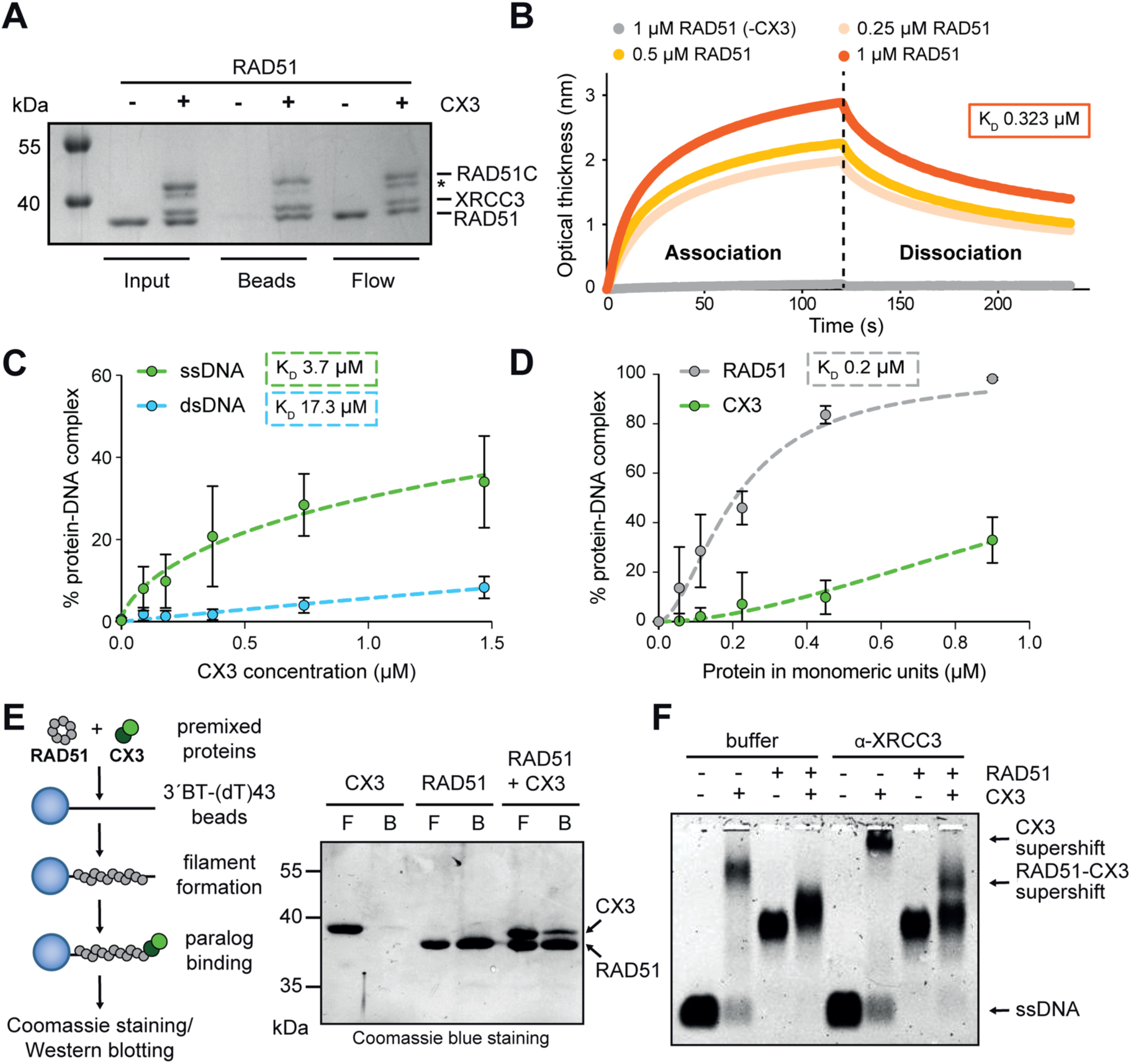
Purified human CX3 complex interacts with RAD51 and RAD51 filaments. **(A)** Pull-down assay with purified RAD51 and His_6_-RAD51C-XRCC3 (CX3) using Ni-NTA beads. Asterisk represents non-specific band. **(B)** Bio-layer interferometry (BLI) sensorgram of the CX3-RAD51 interaction. His-tagged CX3 (0.5 µM) was immobilised on NTA-biosensors and increasing concentrations of RAD51 were bound in the presence of ATP/Mg^2+^. Association and dissociation phases were monitored in real time. The dissociation constant is shown in the orange rectangle. **(C)** Quantification of CX3 binding to ssDNA (green) and dsDNA (blue) from Fig. S2A. CX3 (0.09-1.470 µM) was incubated with 30 nM 49mer ssDNA or dsDNA, crosslinked with 0.5 µL of 0.2% glutaraldehyde, and resolved on 0.8% TBE agarose gels. DNA bounding (in %) was quantified. Error bars indicate ± s.d. (n = 3). Curves represent four-parameter non-linear regression fits. The dissociation constants are shown in the rectangles. (**D**) Quantification of ssDNA binding by RAD51 (0.056-0.9 µM, blue) or CX3 (0.056-0.9 µM, orange) to 30 nM 61mer ssDNA as shown in Fig. S2B. DNA binding (in %) was quantified as in (C). Error bars indicate ± s.d. (n = 3). Curves represent four-parameter non-linear regression fits. The dissociation constant is shown in the orange rectangle. (**E**) Schematics of capture assay with purified proteins (left). Representative Coomassie blue-stained SDS-PAGE of the assay (right), where CX3 (0.94 μM), RAD51 (6.5 μM), or both together were incubated with 3’ biotinylated (BT)-dT43mer ssDNA immobilised on streptavidin-coated magnetic beads for 10 min at 37°C. The flow-through (F) was collected, the beads were washed twice, and bound proteins were eluted (B). (**F**) Representative EMSA demonstrating interaction of CX3 with RAD51 nucleoprotein filament. RAD51 (0.6 μM) filament was assembled on 10 nM 61mer ssDNA in the presence or absence of CX3 (0.6 μM). After glutaraldehyde crosslinking, protein-DNA complexes (PCs) were incubated with an α-XRCC3 antibody to generate a supershifted species. Protein-DNA complexes were resolved on 0.8% TBE agarose gels.

Next, we examined the DNA binding properties of CX3 complex. Electrophoretic mobility shift assays (EMSAs) showed that CX3 forms discrete protein-DNA complexes with a preference for ssDNA (*K*_D_ of 3.7 μM) over dsDNA (*K*_D_ of 17.3 μM) substrates (**Fig. 1C and S2A**). However, compared with RAD51 (*K*_D_ of 200 nM), CX3 alone displayed markedly weaker binding to ssDNA (**Fig. 1D and S2B**). Despite the limited intrinsic DNA-binding activity, CX3 complex was efficiently recovered in capture assays when premixed with RAD51 and assembled on biotinylated ssDNA, irrespective of whether the biotin was positioned at the 5′ or 3′ end (**Fig. 1E and S2C, D**). Consistently, EMSA experiments revealed that addition of CX3 leads to a RAD51 nucleofilament species with reduced electrophoretic mobility (**Fig. 1F and S2E**). This species could be further super-shifted with an antibody against XRCC3 (**Fig. 1F**), confirming the presence of CX3 within the complex. Together, these data demonstrate that CX3 forms a stable complex with RAD51 and preferentially associates with RAD51-ssDNA filaments rather than binding DNA independently.

### CX3 caps and stabilises RAD51 5’ filament ends

To further investigate how CX3 engages RAD51 filament, we first analysed pre-assembled RAD51 filaments by negative-stain electron microscopy (EM). In addition to individual RAD51 filaments assembled on 90mer ssDNA (∼50 nm in length), approximately 30% of the filaments formed longer structures emerging from their end-to-end stacking (**Fig. 2A and S3A**). Increasing concentrations of CX3 progressively reduced the frequency of these stacked filaments, with only ∼3% remaining at a RAD51 ratio of 1:4 (**Fig. 2A and S3A**). This concentration-dependent suppression of filament stacking suggested that CX3 associates with the filament termini and prevents end-to-end stacking. To directly localise CX3 on RAD51 filaments, we carried out immunogold EM with gold-labelled anti-XRCC3 antibodies. Incubation of CX3 with RAD51 led to marked increase in gold-particle association with filament compared to RAD51 alone, with signal enrichment notably at filament ends (9-fold increase) and, to a lesser extent, distributed along the filament (2.6-fold increase) (**Fig. 2B**), consistent with earlier observations in *C. elegans*^34^. As expected, no filament-like structures were formed in the absence of DNA (**Fig. S3C**). Combined with limited intrinsic ssDNA binding of CX3 observed above, these data indicate that CX3 does not specifically recognise ssDNA termini but instead preferentially caps RAD51 filaments through protein-protein contacts at filament ends.

**Fig. 2:**
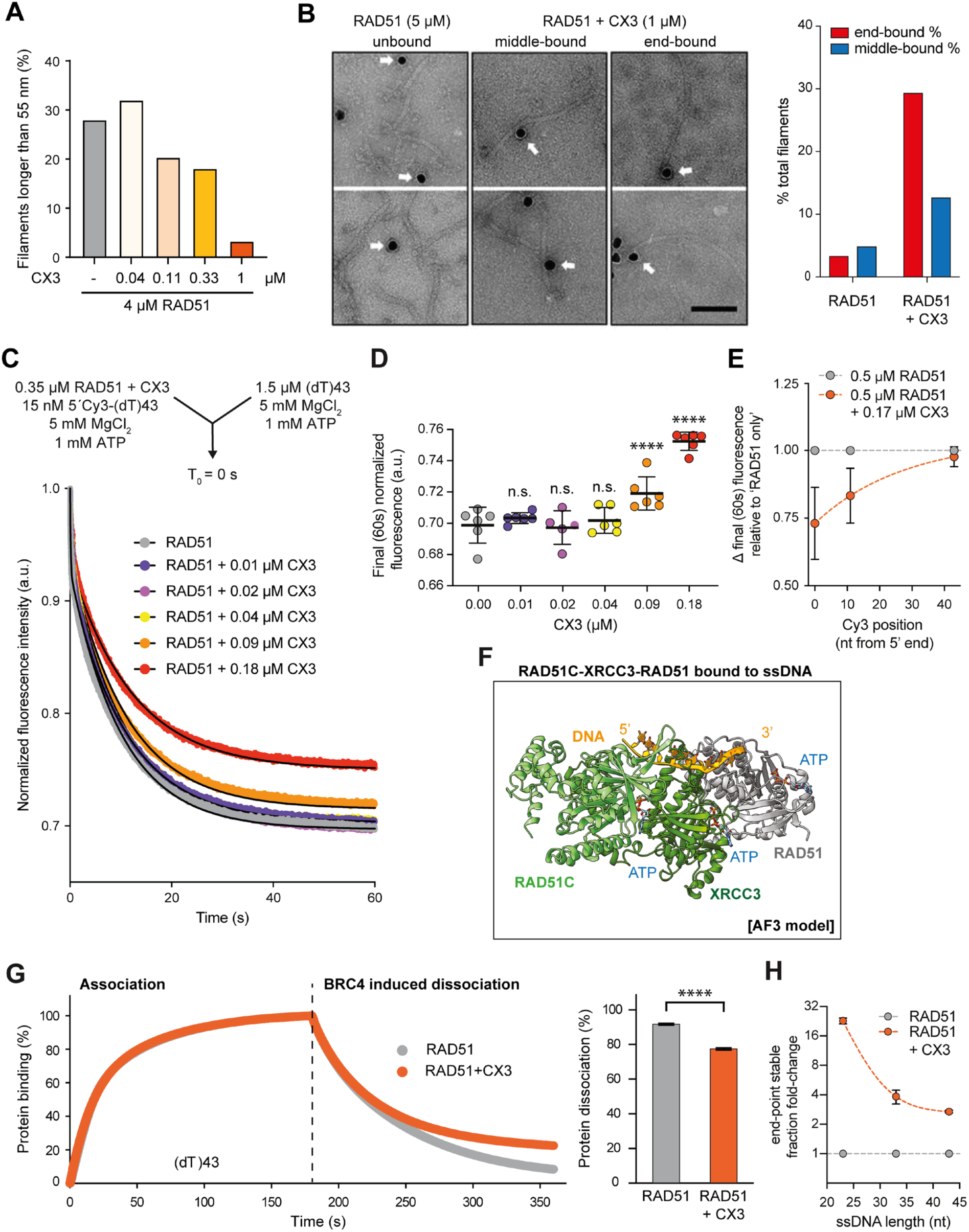
The CX3 preferentially recognises and stabilises 5’ ends of RAD51 filament. **(A)** Percentage of RAD51 filaments longer than 55 nm formed on 90mer ssDNA in the absence or presence of CX3 (0.04–1 µM), based on the negative-stain EM micrographs shown in Fig. S3A and the analysis shown in Fig. S3B.**(B)** Representative negative-stain immunogold EM images (left) of RAD51 filaments assembled on 150mer ssDNA substrate and incubated with CX3. CX3 was detected using α-XRCC3/α-mouse-20-nm gold-conjugated antibodies. Scale bar,100 nm (right). Percentage of RAD51 filaments containing 20-nm gold particles at filament ends (end-bound) or along the filament (middle-bound) in the absence or presence of CX3 respectively. **(C)** Stop-flow analysis of CX3-mediated stabilisation of RAD51 filaments. Schematics of the reaction (top): RAD51 (0.35 µM) was pre-incubated with 15 nM 5′Cy3-(dT)43 for 10 min, in the absence or presence of increasing concentrations of CX3, rapidly mixed with excess of unlabelled (dT)43 ssDNA, and fluorescence was monitored. Average normalised Cy3 fluorescence was plotted as a function of time (bottom). Black lines represent double-exponential fit. (**D**) Quantification of normalised endpoint (60s) Cy3 fluorescence for the data presented in (C). Error bars indicate ± s.d. (n = 4-7). Statistical significance was determined using two-tailed Student’s t test: n.s. not significant, **** p < 0.0001. **(E)** Relative normalised end-point (60s) Cy3 florescence in stopped-flow assays following competitor challenge of RAD51 or RAD51-CX3 filaments assembled on (dT)43 ssDNA labelled with Cy3 at 5’ end (0-nt), internally (11-nt), or at the 3’end (43-nt). Data correspond to average fluorescence traces presented in Fig. S4D. Error bars indicate ± s.d. (n = 4-7). **(F)** AlphaFold 3 predicted structure of ATP-bound human CX3-RAD51 complex formed on 20mer ssDNA. **(G)** Bio-layer interferometry (BLI) sensorgram of RAD51 filament stability (left). Binding of RAD51 (0.75 μM), with (orange) or without (grey) CX3 (0.75 μM) in presence of ATP/Mg^2+^ to 15 nM 3’ biotinylated ssDNA (dT)43 immobilised to streptavidin biosensors was monitored. Filament dissociation was facilitated by addition of 10 μM BRC4 peptide. Average data of 3 independent experiments. Quantification of the endpoint normalised optical thickness from BLI experiment (right). Error bars indicate ± s.d. (n = 3). Statistical significance was determined using two-tailed Student’s t test: **** p < 0.0001. **(H)** Relative fold-change of normalized dissociation end-point optical thickness signal between RAD51+CX3 and RAD51 only condition, plotted as a function of poly(dT) oligonucleotide length. Orange dashed line represent single-exponential fit. Related to data in Fig. S4E.

Given that capping the end of a RAD51 filament may influence its stability, we next assessed the effect of CX3 on RAD51 dissociation form DNA. To validate this in real time, we conducted complementary stopped-flow assays. First, assembly of RAD51 filaments on 5’-Cy3-labelled ssDNA results in an increase in Cy3 fluorescence (**Fig. S4A**), due to a phenomenon referred to as Protein-Induced Fluorescence Enhancement (PIFE). Subsequent challenge with excess unlabelled competitor DNA (cDNA) leads to time-dependent decrease in fluorescence as RAD51 dissociates (**Fig. S4C**). Filament stability was quantified as the fluorescence signal at 60 s relative to initial signal recorded immediately after mixing (∼0.01 s). Increasing CX3 concentrations suppressed the fluorescence decrease in a concentration-dependent manner (**Fig. 2C, D**). Importantly, CX3 alone had no detectable DNA-binding activity under these conditions (**Fig. S4B**), confirming that RAD51 filament stabilisation arises from the CX3-RAD51 interaction rather than from direct DNA binding.

To identify which filament end is being recognized and stabilized by CX3, we monitored the CX3-mediated stabilization of RAD51 filaments assembled on ssDNA labelled with Cy3 at 5’-terminus, 3’-terminus or internally. CX3-induced stabilisation was observed when the Cy3 fluorophore was positioned at the 5’ end or the internal site, but not at the 3’ end of ssDNA (**Fig. 2E and S4D**), suggesting that CX3 shows polarity towards the 5’ end of RAD51 filament in a manner analogous to nematode RAD-51 filament stabilization by RFS-1/RIP-1 complex^32^. Consistently, AlphaFold3^36^ prediction of RAD51C-XRCC3-RAD51-ssDNA complex (ipTM = 0.78) supports preferential engagement of CX3 with the 5’ end of RAD51 filament, with XRCC3 forming the principal interface with the terminal RAD51 protomer (**Fig. 2F**). This arrangement resembles the previously predicted structure of budding yeast Rad55-Rad57-Rad51 complex^37^ and is consistent with recent experimental cryo-EM structures showing X3CD2 complex capping the 5’end of RAD51 filament^12,13^. Importantly, it also parallels our previous findings that the *C. elegans* RFS-1/RIP-1 complex dynamically engages the 5′ end of RAD-51 filaments to promote their stabilisation and remodelling^33,34^. Together, these structural and biochemical observations support an evolutionarily conserved mechanisms of RAD51 filament 5’ end capping by RAD51 paralogs.

Next, we used BLI to test whether CX3 could also stabilise and protect RAD51 filaments from active disassembly by the BRC4 peptide, a BRCA2-derived motif known to disrupt RAD51 filaments^38^. RAD51 filaments were assembled on ssDNA-conjugated biosensors and challenged with the BRC4 peptide. While BRC4 alone efficiently dissociated RAD51 from ssDNA, the dissociation was suppressed in the presence of CX3 (**Fig. 2G**), demonstrating a protective effect of CX3 on RAD51 filaments against both passive competitor-driven turnover and active disassembly by BRC repeats. Because RAD51 filament stability is inherently reduced on shorter ssDNA substrates, we extended the assay to 33mer and 23mer poly(dT) ssDNA. The stabilising effect of CX3 inversely correlated with the ssDNA length: from ∼2.5-fold stabilization on the long (dT)43 ssDNA, up to ∼20-fold stabilization on the short (dT)23 ssDNA (**Fig. 2H and S4E**), suggesting that CX3-mediated capping may be particularly important when filament assembly is kinetically or structurally constrained.

### CX3 remodels RAD51 filament ends into a DNase I-sensitive state

To test the functional consequences of CX3-mediated stabilisation, we employed DNase I protection assays^34^ to probe DNA accessibility within the RAD51 filament. While DNase I efficiently digested naked ssDNA, RAD51-bound ssDNA was substantially protected from degradation (**Fig. 3A, B**). Strikingly, addition of CX3 produced a concentration-dependent increase in DNase I sensitivity, while CX3 alone had no effect on DNase I activity **(Fig. 3A, B**). Since this change was not accompanied by filament disassembly (**Fig. 2C, D**), it cannot be attributed to RAD51 dissociation but rather may indicate that filament end engagement by CX3 gives rise to a compositionally or structurally distinct state of the CX3-RAD51 co-filament in which the bound ssDNA is more accessible to nuclease cleavage while overall filament integrity is preserved.

**Fig. 3:**
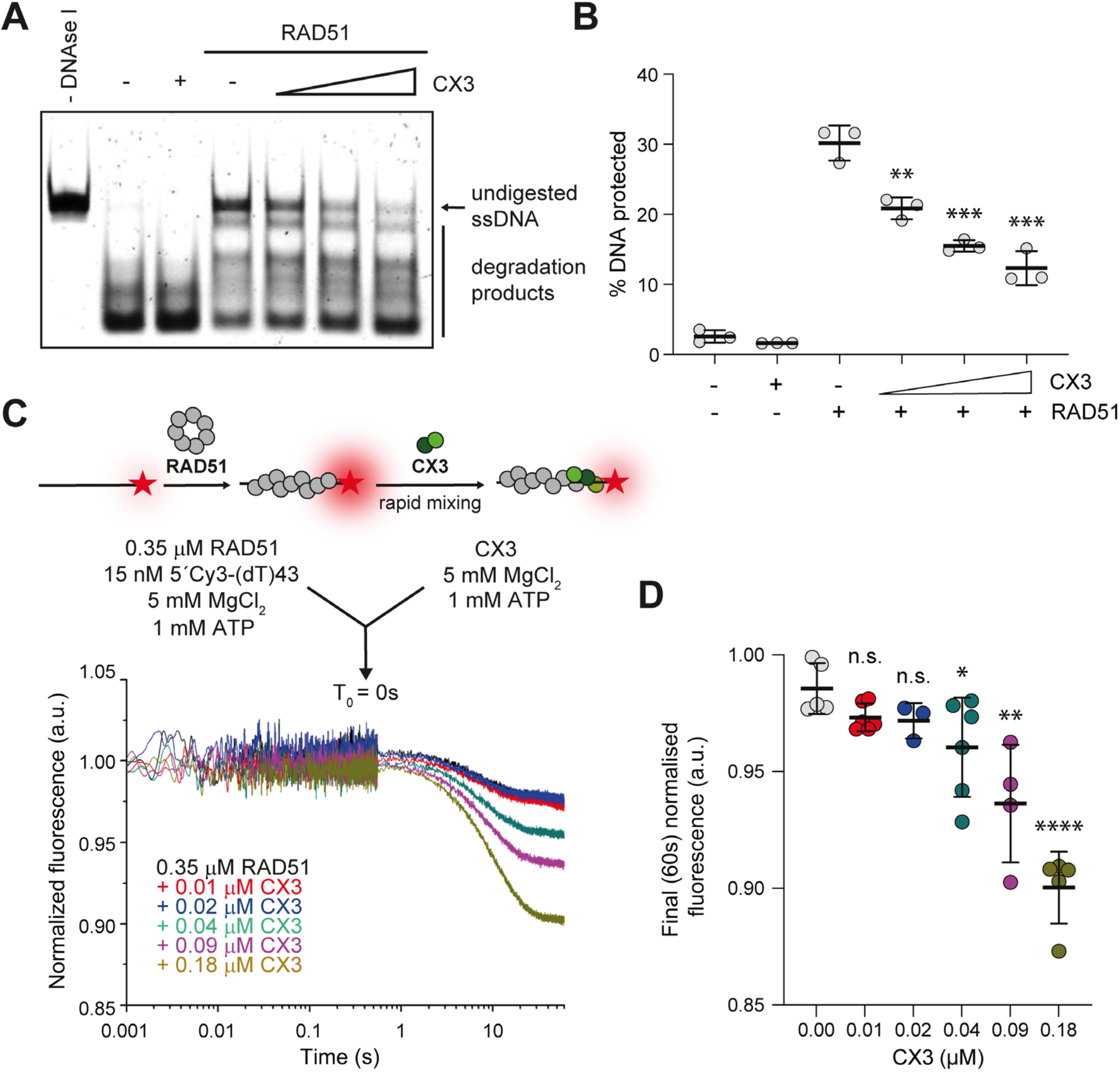
CX3 remodels RAD51 filament ends into a distinct nucleoprotein state. **(A)** Representative PAGE gel of DNase I protection assay. RAD51 (0.6 µM) was incubated with 30 nM 61mer ssDNA in the presence or absence of CX3 (0.15, 0.3, 0.6 µM) for 10 min at 37°C. Protein-DNA complexes were challenged with 1 U DNase I for 20 min at 37°C. Reactions were terminated using 0.125% SDS and 15.6 μg proteinase K. **(B)** Quantification of protection assay shown in panel (A). Error bars indicate ± s.d. (n = 3). Statistical significance was determined using two-tailed Student’s t test: ** p < 0.01, *** p<0.001 **(C)** Schematic of stopped-flow assay monitoring CX3-mediated remodelling of pre-assembled RAD51 filament (top). RAD51 (0.35 µM) was pre-incubated with 1 mM ATP/Mg^2+^ for 5 min and then assembled on 15 nM 5′Cy3 labelled (dT)43 ssDNA for further 5 min. Pre-formed RAD51 filaments were rapidly mixed with CX3 (0.01-0.18 µM) in stopped-flow apparatus and fluorescence was monitored. Average normalised 5′-Cy3-(dT)43 fluorescence is plotted as a function of log_10_(time) (bottom). **(D)** Quantification of normalised endpoint (60s) Cy3 fluorescence for the data presented in (C). Error bars indicate ± s.d. (n = 3-6). Statistical significance was determined using two-tailed Student’s t test: n.s. not significant, * p < 0.05, ** p < 0.01, **** p<0.0001.

To further characterise this structural transition, we performed stopped-flow experiments monitoring fluorescence changes of pre-formed RAD51 filaments in real time following CX3 addition (**Fig. 3C**). We observed a CX3 concentration-dependent decrease in 5’-Cy3 fluorescence (**Fig. 3C, D**), which stems from a protein binding mode change in the vicinity of the 5’-Cy3 dye. This observation is consistent with CX3 mediated induction of a more ‘open’ conformation of 5’ RAD51 filament ends. This observation is consistent with CX3 inducing local rearrangements of RAD51-DNA contacts rather than filament dissociation. Notably, a similar decrease in 5′-Cy3 fluorescence was previously observed upon addition of the nematode RAD51 paralogue complex RFS-1/RIP-1 to pre-assembled RAD-51 filaments, where the fluorescence change extended approximately 6-7 nucleotides from the 5′ ssDNA end^33^. Together, these findings suggest that CX3 remodels the terminal region of the RAD51 filament, generating a stable but conformationally distinct nucleoprotein complex with increased accessibility of the bound DNA.

### CX3 stabilises RAD51 filaments independently of RAD51 ATPase modulation

ATP binding and hydrolysis by the Walker A and B boxes plays a major role in recombinase filament turnover and function. Previous work has established that RAD51 filament stability is enhanced by conditions that impair ATP hydrolysis, including the ATPase-deficient K133R mutation, substitution of Mg^2+^ with Ca^2+^, non-hydrolysable ATP analogues, and engagement by the BRC repeats of BRCA2^39–42^. Given that both subunits of the CX3 complex as well as RAD51 itself contain conserved nucleotide-binding Walker A and B boxes, we aimed to examine the role of ATP in CX3 function.

CX3 itself displayed only minimal intrinsic ATPase activity, irrespective of the presence of ssDNA (k_cat_^CX3^ = 0.05 ± 0.01 min^−1^; k_cat_^CX3+ssDNA^ = 0.04 ± 0.02 min^−1^) (**Fig. S5A**). Furthermore, CX3 addition had no measurable effect on RAD51 ATPase activity (k_cat_^RAD51^ = 0.34 ± 0.04 min^−1^; k_cat_^RAD51(CX3–adj)^ = 0.31 ± 0.02 min^−1^) (**Fig. 4A**), indicating that filament stabilisation by CX3 is not achieved through global modulation of RAD51 ATP turnover, but is mechanistically distinct from the mode of action of BRCA2 BRC repeats^43^.

**Fig. 4:**
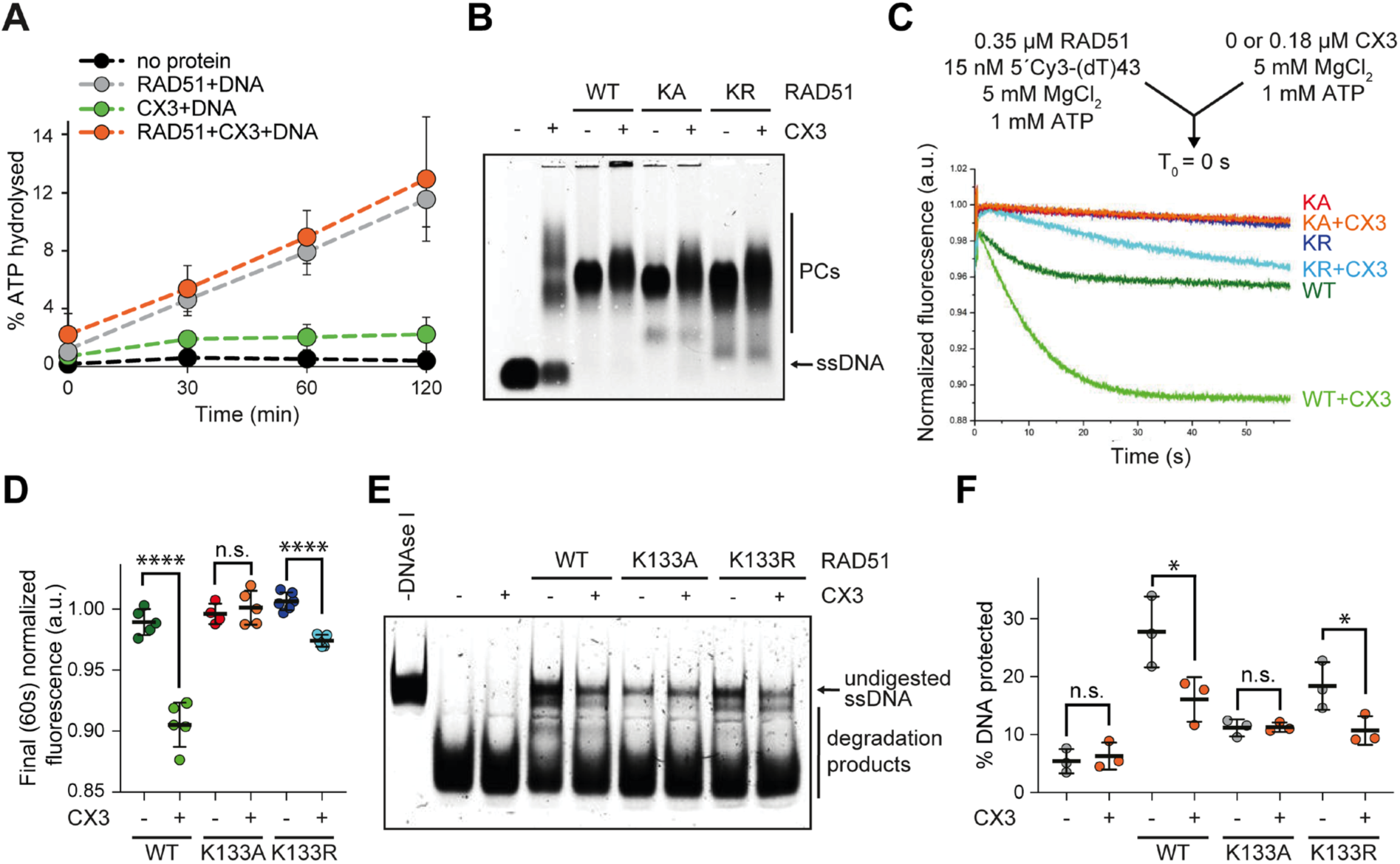
CX3 selectively remodels ATP-bound RAD51 filaments. **(A)** ATPase assay performed with CX3 (0.25 μM) and/or RAD51 (0.25 μM) in the presence of 40 nM (dT)63 ssDNA. Error bars indicate ± s.d. (n = 3). (**B**) Representative EMSA of RAD51 WT and Walker A mutants K133A (KA) or K133R (KR) (1.2 μM each) binding to 30 nM 61mer ssDNA in the presence or absence of CX3 (0.3 μM). Protein-DNA complexes (PCs) were stabilised by crosslinking with 0.5 µL of 0.03% glutaraldehyde and resolved on 0.8% TBE agarose gels. (**C**) Schematics of stopped flow analysis of CX3-mediated remodelling of RAD51 WT and Walker A mutant filaments. The arrow indicates the components of the two syringes rapidly mixed at the ‘0 s’ time point in a stopped-flow instrument (top). RAD51 WT, K133A, or K133R (0.35 μM each) were pre-incubated with 15 nM 5′Cy3-(dT)43 for 10 min, followed by rapid mixing with CX3 (0.18 μM) and monitoring of Cy3 fluorescence. Average normalized 5′Cy3-(dT)43 fluorescence is plotted as a function of time (bottom). (**D**) Quantification of normalised endpoint (60s) Cy3 fluorescence shown in (C). Error bars indicate ± s.d. (n = 4-6). Statistical significance was determined using two-tailed Student’s t test: n.s. not significant, **** p < 0.0001. (**E**) Representative image of DNase I protection assay. RAD51 WT, K133A, or K133R (0.23 μM each) were incubated with 30 nM 61mer ssDNA in the presence or absence of CX3 (0.23 μM) for 10 min. No protein and CX3 alone reactions were used as controls. Protein-DNA complexes were challenged with 1 U DNase I for 20 min. Reactions were terminated using 0.125% SDS and 15.6 μg proteinase K and resolved in 10% native polyacrylamide gels. **(F)** Quantification of protection assay shown in panel (E). Error bars indicate s.d. (n = 3). Statistical significance was determined using two-tailed Student’s t test: n.s. not significant, * p < 0.05.

### The nucleotide state of RAD51 determines CX3 engagement

To dissect the nucleotide requirements of the CX3-RAD51 interaction, we employed two RAD51 Walker A box mutants: RAD51 K133A (KA) and RAD51 K133R (KR), defective in ATP binding and hydrolysis, respectively^40^. BLI analysis revealed that CX3 binds RAD51 wild-type (WT) and both RAD51 mutants (**Fig. S5B, C**). The ATP-binding-deficient KA mutant bound CX3 with reduced affinity (*K*_D_= 563 nM), whereas the hydrolysis-deficient KR mutant showed even tighter binding (*K*_D_= 270 nM) than RAD51 WT (*K*_D_= 323 nM), possibly reflecting preferential recognition of an ATP-bound, non-hydrolytic RAD51 state.

EMSA confirmed association of CX3 with RAD51 filaments formed by WT, KA, and KR (**Fig. 4B**), and the stopped-flow assay confirmed that CX3 stabilises all three filaments at the 5’ fluorescently labelled filament end, even though the KR filaments already displayed very high intrinsic stability (**Fig. S5D-G**), limiting the stabilizing effect dynamic range of the stabilizing effect. Given that CX3 does not modulate RAD51 ATP turnover (**Fig. 4A**), these data indicate that CX3 recognizes 5’ filament ends beyond the state resembling the non-hydrolysable RAD51-ATP conformation.

In contrast to filament binding, the filament end-remodeling activity of CX3 was strictly ATP-dependent. The CX3-induced 5’-Cy3 fluorescence decrease (**Fig. 4C, D**), our proxy for 5’ filament end remodeling, and the transition to the DNase I-sensitive state (**Fig. 4E, F**) were both observed with RAD51 WT and KR, but not with the KA filaments. Both mutant filaments displayed higher baseline DNase I sensitivity than WT, consistent with their reduced ssDNA binding (**Fig. S5H**). In agreement with a defect in remodeling, CX3-mediated RAD51 filament stabilization (measured as increased final (60 s) Cy3 fluorescence upon mixing with ssDNA competitor in stopped-flow setup) failed to propagate at internal or 3′ positions of (dT)43 ssDNA in KA filaments and instead caused a modest destabilization in comparison to WT (**Fig. S5I, J**). These observations indicate that although CX3 can engage KA filaments, it does so in an ineffective manner and cannot trigger the structural rearrangement required for 5’ filament end remodeling and extensive filament stabilization, which propagates towards the middle of the filament. Strikingly, incubation with RAD51 KA, but not with WT or KR, induced robust CX3 ATPase activity (k_cat_^CX3(RAD51 KA-adj)^ = 0.30 ± 0.03 min^−1^) (**Fig. S5K, L**). Because RAD51 KA is itself ATPase-deficient (k_cat_^(RAD51 KA)^ = 0.07 ± 0.03 min^−1^), this activity must arise from CX3, suggesting that the complex engages in a futile ATPase cycle when it binds and attempts to properly engage and remodel a 5’ end of an ATP-free RAD51 filament. Collectively, while CX3 can bind RAD51 filaments independently of RAD51 nucleotide state, the proper functional engagement and 5’ filament end remodeling requires ATP-bound RAD51.

### 5’ RAD51 filament end remodeling requires ATP hydrolysis

We next assessed how the cofactor present in the reaction modulates CX3 function. EMSA demonstrated a more pronounced mobility shift of RAD51 filaments upon CX3 addition with AMP-PNP than with ATP (**Fig. 5A**), suggesting altered filament dynamics in the non-hydrolytic state. With ADP, by contrast, CX3 formed large, aggregated complexes with ssDNA that failed to enter the gel and addition of RAD51 partially rescued entry of CX3 complexes into gel, presumably through formation of more stable co-complexes (**Fig. 5A**). Crucially, CX3-induced DNase I sensitisation was observed only with ATP, and not with AMP-PNP or ADP (**Fig. 5B, C**), indicating that 5’ filament end remodeling requires ATP hydrolysis in the system. To exclude the possibility that the absence of DNAse I sensitisation simply stems from high stability of RAD51 filaments in the presence of AMP-PNP, these experiments were repeated at lower RAD51 concentrations, where comparable results were observed for both AMP-PNP and ATP-γ-S, a second non-hydrolysable analogue (**Fig. S6A-D**). CX3 alone did not protect ssDNA under any nucleotide condition (**Fig. S6E**), confirming the RAD51-dependence of the effect. Together, these findings suggest that ATP binding is sufficient to support CX3 association with RAD51 filaments, whereas ATP hydrolysis is specifically required for the remodeling transition.

**Fig. 5:**
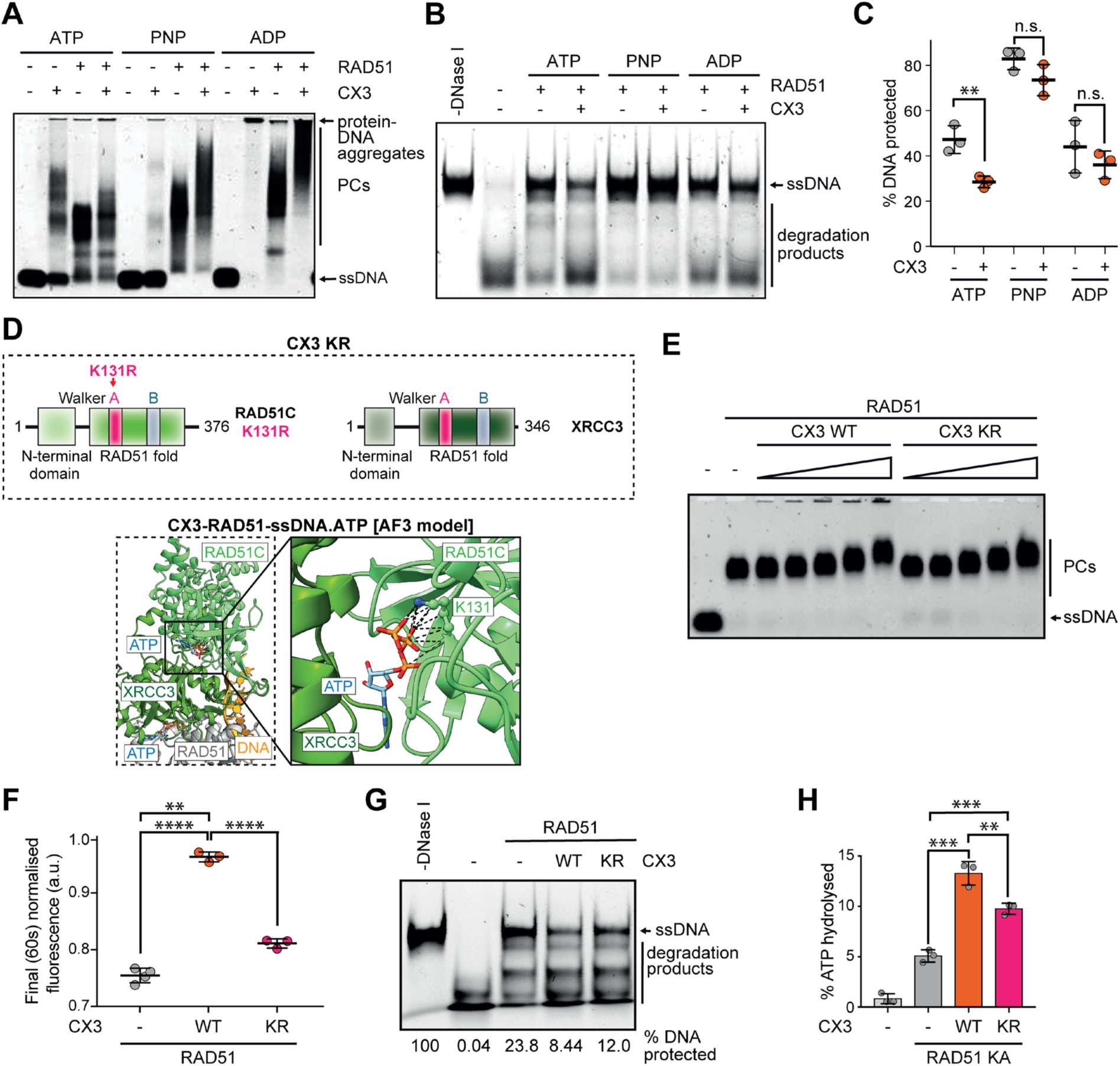
ATP hydrolysis regulates CX3 turnover and remodeling of RAD51 filament ends. **(A)** Representative EMSA of RAD51 and/or CX3 (0.6 μM each) binding to 30 nM 61mer ssDNA in the presence of indicated nucleotide cofactors. Protein-DNA complexes (PCs) were stabilised by crosslinking with 0.5 µL of 0.03% glutaraldehyde and resolved on 0.8% TBE agarose gels. **(B)** Representative image of DNase I protection assay. RAD51 (0.23 μM) and/or CX3 (0.23 μM) were incubated with 30 nM 61mer ssDNA in the presence of indicated nucleotide cofactors for 10 min. Protein-DNA complexes were challenged with 1 U DNase I for 20 min. Reactions were terminated using 0.125% SDS and 15.6 μg proteinase K and resolved in 10% native polyacrylamide gels. **(C)** Quantification of protection assay shown in panel (B). Error bars indicate ± s.d. (n = 3). Statistical significance was determined using two-tailed Student’s t test: n.s. not significant, ** p < 0.01. **(D)** Schematic of RAD51C and XRCC3 proteins. Engineered RAD51C Walker A mutant (K131R) is highlighted in pink. The corresponding complex being referred to as CX3 KR (top). AlphaFold3 predicted structure of ATP-bound CX3-RAD51-ssDNA complex (bottom). RAD51C Walker A lysine and its interaction with the ATP molecule was highlighted in detail. **(E)** Representative EMSA image comparing the effect of CX3 WT and CX3 KR on RAD51 filaments. RAD51 (0.6 µM) was bound to 30 nM 61mer ssDNA in the absence or presence of increasing concentrations (0.037-0.60 µM) of either CX3 WT or CX3 KR. Protein-DNA complexes (PCs) were stabilised by crosslinking with 0.5 µL of 0.2% glutaraldehyde and resolved on 0.8% TBE agarose gels. **(F)** Normalised endpoint (60s) Cy3 fluorescence of RAD51 filaments assembled in the presence of CX3 WT or CX KR following rapid mixing with excess unlabelled competitor ssDNA. Quantification corresponds to the data presented in Fig. S6F. Error bars indicate ± s.d. (n = 3-4). Statistical significance was determined using two-tailed Student’s t test: ** p < 0.01, **** p < 0.0001. **(G)** Representative DNase I protection assay comparing the effect of CX3 WT and CX KR on RAD51 filaments. RAD51 (1.0 μM) was incubated with 30 nM 61mer ssDNA in the presence or absence of CX3 wt or CX3 KR (0.25 μM) for 10 min before challenge with 1 U DNase I for 20 min. Reactions were terminated using 0.125% SDS and 15.6 μg proteinase K and in 10% native polyacrylamide gels. Representative of 3 independent experiments obtained with similar results. Quantification of ssDNA protection in the representative gel is shown below the gel image. **(H)** Quantification of fraction of ATP hydrolysed after 1 h in ATPase reactions containing CX3 WT or CX3 KR (0.4 µM each) in the presence of RAD51 K133A (1 μM) and 160 nM (dT)63 ssDNA. Error bars indicate s.d. (n = 3). Statistical significance was determined using two-tailed Student’s t test: ** p < 0.01, *** p<0.001.

### The RAD51C Walker A motif of CX3 drives the remodeling transition

Given the CX3-RAD51 filament assembly performed in presence of non-hydrolysable ATP analogues completely prevented CX3-mediated RAD51 filament remodeling, but ATPase-deficient RAD51 K133R filaments were still remodelled in the presence of ATP and CX3, we hypothesized that ATP hydrolysis within the CX3 complex may drive the RAD51 filament remodeling. To identify which CX3 subunit supplies the hydrolytic activity, we attempted K to R substitutions in the Walker A box of both RAD51C and XRCC3. Combined Walker A mutations or mutation in the XRCC3 Walker A motif yielded insoluble complexes, consistent with earlier reports that nucleotide-binding motifs influence CX3 complex assembly and stability. Notably, XRCC3 K113R was previously reported to impair CX3 complex formation and render XRCC3 predominantly insoluble upon bacterial co-expression^44^. We therefore purified CX3 carrying only the RAD51C K131R substitution (CX3 KR), which remained soluble. Structurally, K131 residue of RAD51C acts as the key lysine finger and makes contacts with the phosphate groups of the ATP molecule bound between the XRCC3 and RAD51C subunits of the CX3 complex^12,23^ (**Fig. 5D**). CX3 KR bound RAD51 filaments comparably to wild-type CX3, indicating that RAD51C ATP hydrolysis is dispensable for filament binding (**Fig. 5E**) consistent with binding assays performed in the presence of AMP-PNP (**Fig. 5A**). However, CX3 KR displayed strongly decreased ability to stabilise RAD51 filaments in stopped-flow assays (**Fig. 5F and S6F**) and reduced induction of the DNase I-sensitive state (**Fig. 5G**), identifying ATP hydrolysis by RAD51C as essential for the full remodeling transition that underpins robust filament stabilisation. Consistently, while CX3 WT engaged in a futile ATPase cycle selectively in the presence of RAD51 KA (**Fig. S5K**), CX3 KR displayed substantially reduced, albeit not completely abolished, ATPase activity (**Fig. 5H**), suggesting that the XRCC3 Walker A motif may also contribute to full enzymatic turnover and remodeling.

### Global kinetic analysis defines a co-loading mechanism for CX3-mediated RAD51 filament assembly

To define the mechanism by which CX3 modulates RAD51 nucleation and filament dynamics on ssDNA, we performed global numerical fitting of stopped-flow fluorescence transients acquired under six complementary experimental configurations: (i) RAD51 titration at a fixed CX3 concentration followed by addition of Cy3-labelled (dT)43 ssDNA (**Fig. S7A**); (ii) RAD51 titration with Cy3-labelled dT43 ssDNA in the absence of CX3 (**Fig. S7B**); (iii) CX3 titration at a fixed RAD51 concentration followed by addition of Cy3-labelled (dT)43 ssDNA (**Fig. S7C**); (iv) CX3 titration to pre-assembled RAD51-Cy3-labelled (dT)43 ssDNA filaments (**Fig. S7D**); (v) competitor-DNA challenge of pre-formed RAD51-Cy3-labelled (dT)43 ssDNA filaments in the presence and absence of CX3 (**Fig. S7E**); and (vi) RAD51 titration with fluorescently-labelled CX3 (**Fig. S7F**).

The globally fitted model with corresponding kinetic parameters (**Fig. S8**) defines two parallel kinetic pathways. In the absence of CX3, RAD51 engages ssDNA through a four-step mechanism comprising cooperative binding composed of two binding events (*k*_1_, *k*_-1_) and (*k*_2_, *k*_-2_), isomerisation to the filament state (*k*_3_, *k*_-3_) and subsequent filament rearrangement (*k*_-4_, *k*₋₄), reproducing the kinetic scheme reported previously^22^ (**Fig. S8C**). In the presence of CX3, RAD51 and CX3 first pre-associate in solution (*k*_5_ = 0.005 ± 0.0002 nM⁻¹ s⁻¹; *k*_-5_ = 2.93 ± 0.04 s⁻¹; *K*_D5_ = 0.59 µM, **Fig. S8A, B**) to form a RAD51-CX3 complex that subsequently engages ssDNA through an analogous but kinetically distinct pathway (*k*_7_–*k*_10_; **Fig. 6 and S8C**). Two principal consequences of CX3 engagement emerged from the global fit. First, CX3 substantially enhanced the effective affinity of RAD51 for ssDNA during the initial encounter step. The apparent dissociation constant of the first binding step decreased from *K*_D1_ = 3.0 µM for RAD51 alone to 0.07 µM for the RAD51-CX3 pathway (∼43-fold enhancement), while the second binding step was strengthened ∼7-fold (*K*_D2_ = 0.56 µM versus 0.085 µM) (**Fig. 6 and S8B**). Second, whereas assembly of RAD51 alone required cooperative binding steps, the RAD51-CX3 complex engaged ssDNA in a non-cooperative manner, indicating that pre-formation of the RAD51-CX3 unit in solution circumvents the cooperative binding step required for nucleation of RAD51 alone (**Fig. 6 and S8C**). Together, these data suggest that CX3 converts cooperative RAD51 nucleation into a direct, non-cooperative co-loading mechanism, which would explain enhanced RAD51 filament assembly on short or sub-saturating ssDNA substrates.

**Fig. 6:**
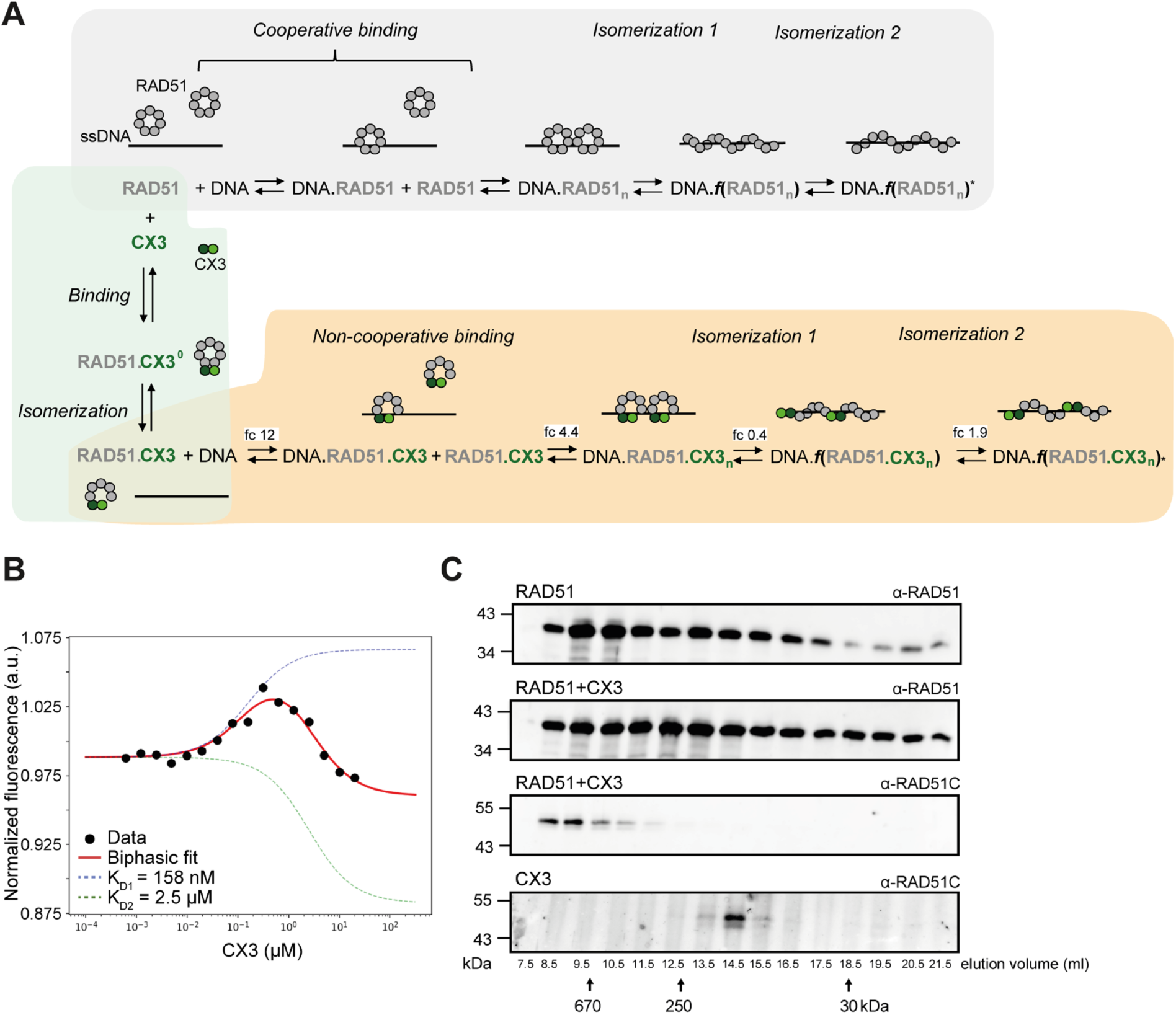
Global kinetic model of RAD51 filament assembly and its regulation by CX3. **(A)** Kinetic model describing RAD51 filament assembly in the absence and presence of CX3, derived from global numerical analysis of stopped-flow fluorescence data. The complete experimental datasets and fitted rate constants are summarized in Fig. S7 and S8, respectively. In the absence of CX3 (grey), RAD51 assembles on ssDNA through cooperative binding followed by two sequential isomerization steps. CX3 interacts with soluble RAD51 through initial binding step followed by isomerisation, generating a CX3-RAD51 complex (green). This complex subsequently engages ssDNA through a non-cooperative binding pathway, followed by two isomerisation steps (orange). Although the oligomeric state of RAD51 during ssDNA loading remains unresolved, RAD51 is schematically represented as a ring for simplicity. This depiction accommodates the potentially heterogeneous and condition-dependent oligomeric states of RAD51 and does not imply a defined ring structure during filament assembly. Fold changes (fc) in the corresponding rate constants for RAD51 assembly in the presence versus absence of CX3 (k_7_/k_+1_, k_8_/k_+2_, k_9_/k_+3_, and k_10_/k_+4_) are indicated, highlighting the effects of CX3 on filament binding, stability and isomerisation dynamics. **(B)** Microscale thermophoresis (MST) analysis of the interaction between fluorescently labelled RAD51 and CX3. Normalised fluorescence is plotted as a function of CX3 concentration. The data display a biphasic binding profile and were fitted using a biphasic binding model (red line), yielding apparent dissociation constants of *K*_D1_ = 158 nM and *K*_D2_ = 2.5 μM. Dashed lines indicate the individual binding components. **(C)** Size-exclusion chromatography analysis of RAD51, CX3, and RAD51 incubated with CX3. Individual fractions were analysed by immunoblotting using anti-RAD51 or anti-RAD51C antibodies, as indicated. Elution volumes are shown below the blots; arrows indicate the elution positions of molecular-mass standards (670, 250, and 30 kDa). Co-incubation with CX3 shifts the RAD51 elution profile toward lower-molecular-mass fractions, consistent with CX3-induced remodelling of RAD51 oligomers and formation of RAD51-CX3 complexes.

The kinetic model further provides a quantitative explanation for the stabilisation and remodeling activities of CX3 on RAD51 filaments. Most notably, CX3 reduces the reverse rate associated with filament dissociation and conformational changes. The reverse rates of the first binding step (*k*_-1_) decreased from 180.0 ± 2.0 s^−1^ to *k*_-7_ = 50.0 ± 2.0 s^−1^ (∼3.6-fold reduction), while the reverse isomerisation rate decreased from *k*_-3_ = 2.09 ± 0.03 s^−1^ to *k*_-9_ = 0.5 ± 0.3 s^−1^ (∼4-fold reduction). These changes are consistent with stopped-flow and BLI data showing that CX3 stabilises the filament against competitor- and BRC4-driven disassembly.

Together, the global kinetic analysis indicates that CX3 acts through two coupled mechanisms: (i) enhancement of RAD51 loading via formation of a pre-assembled RAD51-CX3 complex with increased affinity for ssDNA associated with loss requirement for cooperative binding, and (ii) stabilisation of a remodelled filament state through suppression of key reverse reactions. The kinetic framework thus provides a quantitative explanation for observed stabilisation, remodeling and DNase I-sensitisation activities of CX3 and is compatible with the proposed CX3-RAD51 co-filament model.

### CX3 induces a change in the oligomeric state of RAD51

To investigated structural changes in DNA-free RAD51 state induced by CX3 engagement, we first asked whether CX3 perturbs the oligomeric state of RAD51. Free RAD51 has been reported to adopt a polydisperse distribution of oligomers in solution, including heptameric rings^45,46^. We therefore used AlphaFold3 to model the states of DNA-free RAD51 in the presence (ipTM=0.52) or absence of CX3 (ipTM=0.37). In contrast to RAD51, where the prediction showed a heptameric ring (**Fig. S9A**), replacement of 2 monomers of RAD51 with CX3 resulted in ring opening, distorted the symmetric structure, and made the DNA binding sites more accessible with XRCC3 contacting the terminal RAD51 monomer (**Fig. S9B**).

To experimentally test this possibility, we first performed microscale thermophoresis (MST) using RED-labelled RAD51 incubated with increasing concentrations of CX3. The interaction displayed a biphasic binding profile, consisting of an initial high-affinity binding event (*K*_D1_ ∼158 nM) followed by a second lower-affinity transition (*K*_D2_ ∼2.5 µM) associated with an opposite thermophoretic response (**Fig. 6B**). These data indicate that CX3 binding induces at least two distinct RAD51 states and are consistent with a CX3-induced conformational or oligomeric rearrangement of RAD51. We next analysed the oligomeric distribution of RAD51 and CX3 by size-exclusion chromatography. RAD51 alone eluted predominantly as a high-molecular-weight species (∼700 kDa), consistent with large oligomeric ring assemblies. Incubation with CX3 produced a marked redistribution of the RAD51 elution profile shifted towards smaller mixed assemblies of ∼350 kDa (**Fig. 6C**). In parallel, CX3 shifted from its predominant heterodimeric form (∼100 kDa) into a higher-molecular-weight species (**Fig. 6C**), consistent with its incorporation into mixed RAD51-CX3 oligomers. Together, these data support a model in which CX3 remodels RAD51 oligomeric assemblies and promotes formation of distinct lower-molecular-weight RAD51 complexes.

## DISCUSSION

Recent structural studies have significantly advanced our understanding of mammalian RAD51 paralogs. Cryo-EM analyses revealed that RAD51 paralogs assemble into BCDX2-CX3 and DX2-CX3 supercomplexes, which contact RAD51 at the 5’ filament end, consistent with a capping function^12,13^. While these studies established a structural framework for RAD51 paralog function, the molecular consequences of CX3 engagement with RAD51, particularly the role of ATP binding and hydrolysis in regulating filament architecture and dynamics, remained unclear. By combining biochemistry, real-time biophysics, mutagenesis, and global kinetic analysis, we extend these structural observations showing that CX3 binds RAD51 with nanomolar affinity, promotes filament assembly and preferentially caps the 5′ filament end. CX3 further stabilises RAD51 filament against both passive (competitor DNA) and active (BRC4 peptide) dissociation and actively remodels them into a structurally distinct DNase I-sensitive state. Importantly, we establish that filament stabilisation and remodelling are mechanistically separable processes governed by distinct nucleotide requirements. Whereas CX3 association and initial filament stabilisation depend on the ATP-bound state of RAD51, productive remodelling additionally requires ATP hydrolysis by RAD51C, without altering the intrinsic ATPase activity of RAD51. Thus, our findings extend the structural model of RAD51 paralog-mediated filament capping by identifying ATP hydrolysis by CX3 as the driving force for RAD51 filament remodelling, rather than merely a requirement for filament association or stabilisation.

### CX3 remodels RAD51 oligomeric architecture

Our data suggest that CX3 alters the oligomeric state of RAD51. Free RAD51 has been reported to exist as a polydisperse population of oligomers, including heptameric and higher-order ring structures^47,48^. Our AlphaFold-Multimer predictions indicate that CX3 binding destabilises the closed RAD51 ring architecture, and size-exclusion chromatography revealed a substantial redistribution of RAD51 oligomers into smaller mixed assemblies. MST measurements supported this interpretation, identifying a biphasic CX3-RAD51 interaction in which a high-affinity binding event was followed by a second, lower-affinity transition with an opposite thermophoretic signature. The simplest interpretation is that the initial CX3 binding event is followed by a structural rearrangement of RAD51 oligomers, generating a distinct mixed assembly. Indirect support for this solution-phase rearrangement emerged from our global kinetic analysis, which included a pre-association step between CX3 and RAD51. The ∼16-fold difference in apparent affinities and the opposing thermophoretic responses are consistent with the behaviour of many AAA+ and RecA-family proteins, where ligand binding is coupled to large-scale conformational transitions that alter assembly state and protein-protein interactions^49^. Whether this rearranged state represents open RAD51 rings, mixed CX3-RAD51 oligomers, or an early nucleation intermediate is an important question for future structural studies. These observations align with the recent finding that RAD51 paralog assemblies act as protofilament-like nuclei on ssDNA^12^, suggesting that CX3 may not only stabilise pre-existing RAD51 filaments but also pre-condition unproductive RAD51 oligomers into filament nucleation unit.

### Global kinetic analysis reveals a CX3-mediated co-loading mechanism

Another important functional consequence of CX3-mediated rearrangement of RAD51 oligomeric state emerges from global numerical analysis of stopped-flow fluorescent data obtained across six experimental configurations. The resulting kinetic model identifies two parallel pathways for RAD51 filament assembly. In the absence of CX3, RAD51 assembles on ssDNA through cooperative binding followed by two sequential isomerisation steps, consistent with our previously established four-step scheme mechanism^22^. In contrast, CX3 pre-associates with RAD51 in solution (*K*_d5_ = 0.59 µM), forming a CX3-RAD51 complex that engages ssDNA through a non-cooperative co-loading pathway. This mechanism enhances the effective affinity of RAD51 for ssDNA by ∼43-fold during the initial encounter step and sevenfold during the subsequent binding step, effectively bypassing the kinetic barrier imposed by cooperative RAD51 assembly. These findings suggest a mechanism for the ability of CX3 to promote RAD51 filament formation.

Beyond facilitating filament assembly, the kinetic model reveals how CX3 regulates stability and conformational dynamics. CX3 differentially affects the two sequential isomerisation steps associated with filament maturation. During the first transition, CX3 reduces both forward and reverse rate constants, with a more pronounced suppression of the reverse reaction, thereby favouring the intermediate filament state. In contrast, CX3 accelerates both forward and reverse rate constants of the second isomerisation, indicating enhanced conformational exchange rather than a preferential stabilisation of the final state. Together, these changes modestly favour the final isomerised state relative to the preceding filament state, indicating that CX3 reshapes the kinetic landscape of filament maturation rather than simply accelerating a single conformational transition.

Importantly, these effects on filament dynamics are accompanied by a substantial increase in RAD51-ssDNA affinity and enhanced resistance to dissociation. CX3-mediated stabilisation is particularly pronounced on short ssDNA substrates, suggesting a potential role in maintaining RAD51 filaments on spatially constrained DNA intermediates generated during DSB repair and at stalled replication forks. This mechanism may help explain previous reports linking RAD51 paralogs to replication fork protection and restart^18,23^.

### Comparison with BRCA2 and the BCDX2 paralog complex

Collectively, our data support a multistep model in which CX3 regulates RAD51 filament assembly, stability and maturation through distinct but coordinated activities. CX3 initially interacts with soluble RAD51, altering its oligomeric state and generating an assembly-competent CX3-RAD51 intermediate that engages ssDNA through a non-cooperative co-loading pathway (**Fig. 7A, B**). Following filament assembly, CX3 preferentially engages the 5′ filament end, stabilises the nucleoprotein filament by suppressing dissociation and conformational transitions, and drives ATP-hydrolysis-dependent remodelling into a structurally distinct, DNase I-accessible but kinetically stable co-filament.

**Fig 7.**
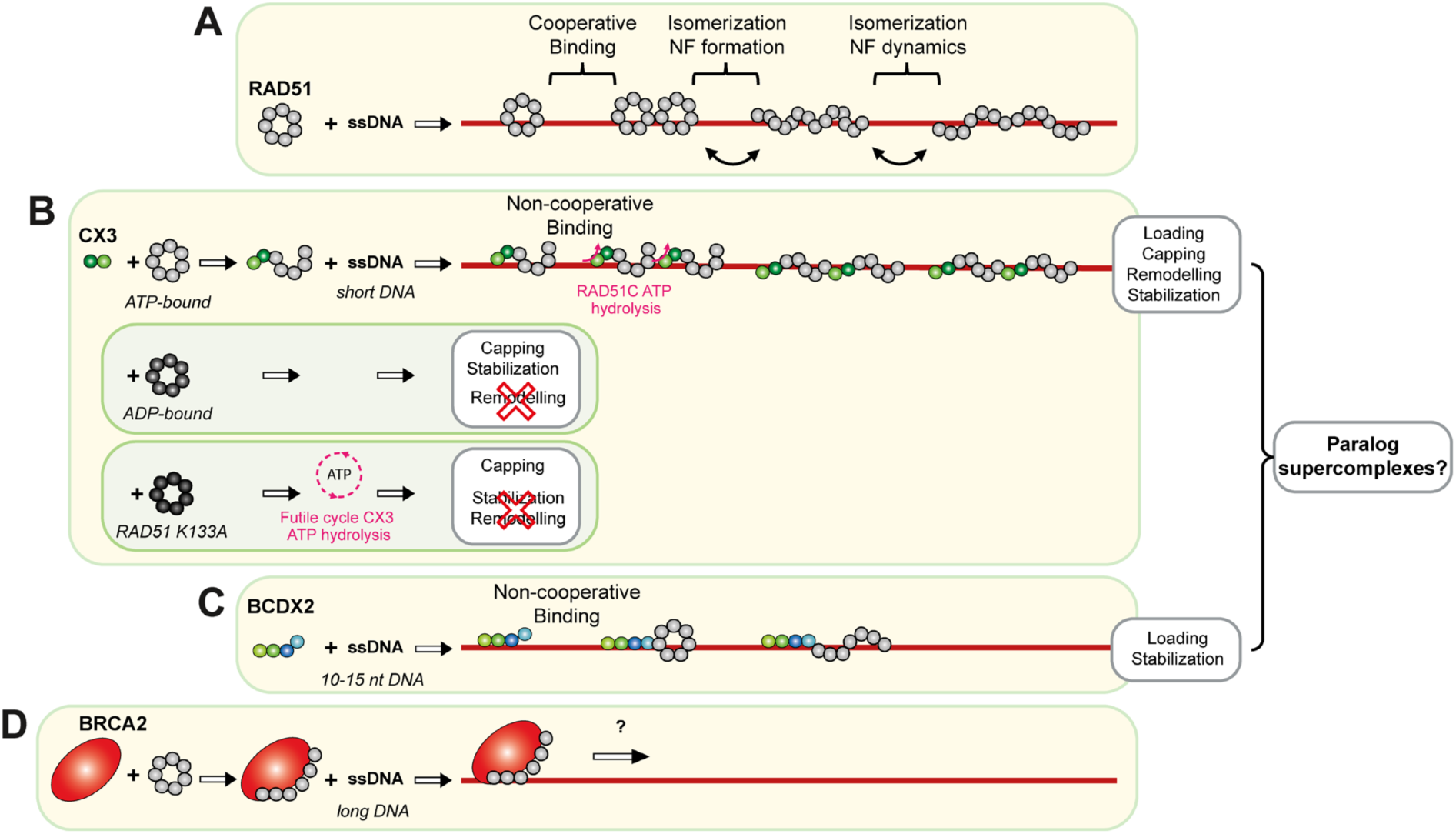
Mechanistic models for RAD51 filament assembly and regulation by RAD51 mediators. **(A)** RAD51 assembles on ssDNA through cooperative binding followed by conformational transitions associated with filament isomerizations. **(B)** Model for CX3-mediated filament assembly and remodelling. CX3 interacts with soluble RAD51, alters its oligomeric state and forms a CX3-RAD51 co-complex that loads onto ssDNA through a non-cooperative pathway, particularly on short DNA substrates. CX3 subsequently caps RAD51 filament ends, promotes filament remodelling and stabilises the resulting nucleoprotein filament. Pink arrows indicate RAD51C-mediated ATP hydrolysis. The effects of ADP-bound RAD51 and RAD51 K133A are shown in the insets. **(C)** Model for BCDX2-mediated RAD51 filament assembly. BCDX2 first binds ssDNA, including short (10-15 nt) substrates, and promotes non-cooperative RAD51 loading and filament formation, followed by filament stabilisation. CX3 and BCDX2 may potentially cooperate within higher-order RAD51 paralog supercomplexes to coordinate filament assembly, capping, remodelling and stabilisation. **(D)** BRCA2 interacts with soluble RAD51 and delivers RAD51 to ssDNA, promoting nucleation and filament assembly on longer ssDNA substrates; subsequent steps and their relationship to RAD51 paralog-mediated filament maturation remain to be defined.

These activities distinguish CX3 from the other major RAD51 mediators, BRCA2 and the BCDX2 complex, while highlighting their complementary role in filament regulation. BCDX2 complex acts through a distinct assembly mechanism, engaging ssDNA first and templating RAD51 nucleation from the DNA-bound state on short ssDNA stretches and RPA-coated substrates^19–22^ (**Fig. 7C**). This DNA-first templating mechanism contrasts directly with the protein-first co-loading pathway identified here for CX3. BRCA2 promotes RAD51 nucleation primarily through its BRC repeats, which sequester RAD51 from non-productive self-association, suppresses RAD51 ATP hydrolysis and facilitate its loading onto RPA-coated ssDNA^9,10,39,43^ (**Fig. 7D**). In contrast, CX3 does not affect the intrinsic RAD51 ATPase activity but instead employs ATP hydrolysis to drive co-filament remodeling.

Thus, although all three mediators facilitate RAD51 filament formation and stability, they achieve these outcomes through different molecular mechanisms. These findings suggest that RAD51 paralogs do not merely reinforce BRCA2-mediated filament assembly but provide additional layers of regulation that control filament architecture and maturation. The mechanistic distinction between CX3 and BRCA2 is further illustrated by the ability of CX3 to counteract BRC4-mediated RAD51 filament disassembly, particularly on short ssDNA substrates. Together with our kinetic analysis, this finding suggests that CX3 stabilises RAD51 filaments by altering their dissociation and conformational dynamics, rather than simply promoting filament assembly. Thus, CX3 not only complements BRCA2-mediated RAD51 loading but also protects assembled filaments against destabilisation by BRC repeats.

Although these mediators promote RAD51 filament assembly and stability, they act through distinct molecular mechanisms. BRCA2 regulates RAD51 availability and nucleotide state to facilitate productive loading, BCDX2 provides an ssDNA-bound platform for filament nucleation, whereas CX3 promotes RAD51 co-loading, preferentially engages the 5′ filament end and mediates ATP-dependent remodelling. An additional mechanism of RAD51 stabilisation has recently been described for the adaptor protein GRB2, which protects RAD51 filaments at reversed replication forks by suppressing RAD51 ATPase activity^50^. In contrast, CX3 does not alter the intrinsic ATPase activity of RAD51 but instead requires an intact RAD51C nucleotide-binding motif to promote filament remodelling into a distinct, DNase I-accessible state. These observations highlight complementary strategies for regulating RAD51 filament stability, ranging from suppression of RAD51 catalytic turnover to active modulation of filament architecture.

Importantly, the recently described BCDX2-CX3 supercomplex architecture^13^ provides a structural basis for potential coordination between these activities. Our findings assign distinct biochemical functions to its constituent modules, suggesting that BCDX2-mediated ssDNA recognition and nucleation could be coupled to CX3-mediated filament stabilisation and ATP-dependent remodelling within a single molecular assembly (**Fig. 7C**). Although the functional coordination of these modules within the intact supercomplex remains to be established, this model provides a framework for understanding how mammalian RAD51 paralogs may integrate filament initiation and maturation to support efficient homologous recombination.

### Conserved 5′-end polarity of RAD51 paralog action

The preferential association of CX3 with the 5′-end of RAD51 filament, supported by immunogold EM, polarity-resolved fluorescence measurements and kinetic analysis, closely resembles the activity of the *C. elegans* RFS-1/RIP-1 complex, which binds 5’ filament termini and promotes maturation through dynamic end engagement^33,34^. This polarity appears to be an evolutionarily conserved feature of RAD51 paralogs, extending from archaeal RadB, capping 5’ ends of RadA filaments^12^, to the yeast Rad55-Rad57 Rad51 paralog dimer ^37,51^ and *C. elegans*^33,52^. These complexes promote Rad51 assembly on RPA-coated ssDNA, stabilise filament ends and facilitate remodelling and subsequent D-loop formation^34,51,52^.

In higher eukaryotes, RAD51 paralogs have diversified into larger assemblies. In species lacking RAD51B, DX2-CX3 complex may provide 5’-end capping activity, consistent with recent structural and biochemical studies^12,13,19^. In mammals, five RAD51 paralogs form two different functional modules: BCDX2 and CX3 both promoting RAD51 nucleation on ssDNA by different molecular mechanisms, with the CX3 complex preferentially engaging the 5’ filament end to stabilise and remodel RAD51 filament. Their integration into a larger supercomplexes may enable tighter coordination of filament assembly and maturation, reflecting the increased regulatory complexity of HR in higher eukaryotes.

More broadly, CX3-mediated filament remodelling shares conceptual similarities with the regulation of ATP-driven filament remodellers, including the actin-severing protein cofilin/ADF^53,54^ and the microtubule-severing AAA+ ATPases katanin and spastin^55,56^, reflecting the deep evolutionary relationship between RAD51 and actin^57,58^.

### ATP coupling and non-productive cycling on RAD51 K133A filaments

Our findings reveal a critical mechanistic distinction between ATP-dependent filament recognition and remodelling. Although CX3 binds RAD51 filaments irrespective of RAD51 nucleotide state, only ATP-bound filaments undergo productive remodeling. Notably, CX3-mediated remodelling does not directly correlate with steady-state ATPase activity. Both WT and K133R RAD51 filaments undergo efficient remodelling despite stimulating little detectable CX3 ATPase activity. Conversely, ATP-binding-deficient RAD51 K133A filaments retain CX3-binding competence and strongly stimulate CX3 ATP hydrolysis but fail to support productive remodelling. These observations indicate that CX3-mediated remodelling depends primarily on the nucleotide-dependent conformational state of RAD51 rather than the overall rate of ATP turnover.

ATP hydrolysis by the paralogs was proposed to act as a release mechanisms^52^ and a structural switch for the active filament state^13^. Our results extend these models by suggesting that ATP turnover is directly coupled to productive filament remodeling. When RAD51 cannot adopt the required ATP-bound conformation, as for the K133A mutant, CX3 binding fails to produce the corresponding structural transition, resulting instead in repeated, non-productive ATP hydrolysis. The reduced affinity of CX3 for RAD51 K133A relative to WT and K133R is consistent with less stable engagement of the defective filament. This behaviour suggests a potential ATP-dependent quality-control mechanism, in which CX3 preferentially stabilises and remodels appropriately configured RAD51 filaments while ATP hydrolysis becomes uncoupled from productive conformational changes on defective substrate. Analogous non-productive ATPase cycles have been observed in several AAA+ remodellers and chaperone systems when substrate engagement occurs without productive structural rearrangement^59^.

Importantly, mutational analysis identifies the RAD51C Walker A motif as the key site of ATP-dependent filament remodeling, while XRCC3 may contributes additional residual activity. These findings distinguish the catalytic contributions of the two CX3 subunits and support a model in which ATP hydrolysis by RAD51C drives the transition from a stabilised RAD51 filament to a distinct remodelled state. Rather than serving solely as a nucleotide-dependent binding or release signal, ATP turnover therefore emerges as a central mechanochemical component of CX3-mediated RAD51 filament maturation.

### Clinical implications of the proposed model for CX3 action

The involvement of the RAD51 paralogs in Fanconi anaemia, hereditary breast and ovarian cancer, and replication stress responses highlights the potential clinical relevance of our findings. By identifying an ATP-dependent filament remodeling step that is mechanistically distinct from the BRCA2-mediated RAD51 loading, our study provides a framework for understanding how disease-associated RAD51 paralogs variants may compromise HR. Notably, several RAD51C mutations affect residues involved in ATP binding, including those within or surrounding the Walker A motif^20^. Recent deep-mutational scanning studies have further revealed clustering of function-impairing RAD51C missense mutations in this region^60^, highlighting the importance of nucleotide-dependent regulation for RAD51C function. Our findings suggest that such pathogenic variants may compromise filament remodeling without necessarily abolishing RAD51 binding, as illustrated by the RAD51C K133R mutant examined here. This distinction may be particularly relevant for interpreting pathogenic mechanisms that cannot be explained solely by defects in RAD51 recruitment or filament assembly. Consistent with this model, cancer-associated RAD51C Walker variant A126T was previously shown to impair RAD51 persistence at stalled replication forks and fork protection despite retaining HR proficiency^23^. Whether these defects arise from impaired CX3-mediated filament remodelling remains to be established. Similarly, Walker A mutations in RAD51C and XRCC3 Walker A mutants were shown to abolish CX3-mediated fork restart in cells^24^. More broadly, by proposing ATP-dependent filament remodeling as a function of RAD51 paralogs, our work provides a mechanistic framework for investigating the functional consequences of cancer-associated variants of uncertain significance in both *RAD51C* and *XRCC3*.

In summary, our study reveals a multistep mechanism by which CX3 regulates RAD51 filament assembly and maturation. CX3 interacts with soluble RAD51 to promote its loading onto ssDNA, preferentially engages the 5′ end of assembled filaments, and couples ATP hydrolysis to the remodelling of ATP-bound RAD51 filaments into a distinct, DNase I-accessible yet kinetically stable state. We propose that this ATP-dependent transition generates a filament architecture favourable for subsequent DNA strand invasion. Together, these findings establish a mechanistic framework linking RAD51 paralog-mediated filament assembly, stabilisation and remodelling, and provide new insights into how defects in CX3 function may contribute to impaired homologous recombination, genome instability and cancer predisposition.

## MATERIALS AND METHODS

### RAD51 and RAD51C/XRCC3 expression

RAD51, RAD51 K133R and K133A were purified as described previously^61^. CX3 protein complex was purified as described previously^23^. Shortly, His6-MBP-TEV-XRCC3/His6-TEV-RAD51C complex was expressed using insect expression system and purified by Ni-NTA agarose (Qiagen), eluted fractions were cleaved by His-TEV-Protease. After additional purification by Ni-NTA agarose, the unbound fraction containing CX3 was loaded on Sephacryl 200 column (Fig. S1A). Alternatively, for pull-down experiment we have cloned His6-MBP-Pre-XRCC3/His6-TEV-RAD51C variant and during purification only Precission Protease was used to get XRCC3/His6-TEV-RAD51C protein complex. RAD51 was labelled by Alexa Fluor™ 555 NHS Ester (Succinimidyl Ester) (Thermofisher scientific A20009) according to manufactureŕs instructions.

### Electrophoretic mobility shift assay (EMSA)

Proteins were diluted in Dilution Buffer (25 mM Tris-HCl pH 7.5, 10% (v/v) glycerol, 0.5 mM EDTA) supplemented with 2 mM MgCl_2_ and 2 mM ATP. To start the reaction, proteins were incubated with prepared mastermix −10 mM Tris, pH 7.5, 10 mM KCl, 1 mM DTT, 2 mM MgCl_2_, 2 mM ATP and indicated concentration of either 5′ FITC labelled 61mer (GACGCTGCCGAATTCTACCAGTGCCTTGCTAGGACATCTTTGCCCA-CCTGCAGGTTCACCC) or 5′ FITC labelled 49mer (AGCTACCATGCCTGCACGAATTAAGCAATTCGTAATCATGGTCATAGCT) oligonucleotide. dsDNA substrates were prepared as described previously^62^. Final reaction volume was 10 µL. Reactions were incubated at 37°C for 10 minutes. To crosslink protein-DNA complexes, 0.5 µL of 0.01-0.2 % (concentration indicated in figure legends) glutaraldehyde was added followed by further 10 min incubation. Products were resolved using 0.8 % TBE agarose gels at 4°C for 60 or 90 min (6.5 V/cm). Gels were imaged on a FLA-9000 scanner (Fujifilm) and quantified with Multi Gauge V3.2 (Fujifilm).

RAD51 was diluted in T50 buffer (25 mM Tris-HCl pH 7.5, 50 mM KCl, 10% glycerol, 0.5 mM EDTA, 1 mM DTT, 0.01% NP40), CX3 in 50 mM Tris-HCl pH 7.5, 50 mM KCl, 1 mM MgCl2 and 1 mM ATP. RAD51 (0.25 µM), Alexa Fluor 555-labelled RAD51 (RAD51-555, 0.13 µM) and increasing concentration of CX3 (0.06 µM-2 µM) were pre-incubated for 1 min at RT followed by transfer to 50 mM Tris-HCl pH 7.5, 50 mM KCl, 1 mM MgCl_2_, 1 mM ATP and 1 mM CaCl_2_ and incubated for 10 min at 37°C. The resulting RAD51-CX3 complex was cross-linked with 0.01% glutaraldehyde for 10 min at 37°C and separated on 0.8% agarose gel in 1x TBE + 10 mM KCl at 70 V for 105 min at 4°C. Gels were imaged by Typhoon™ laser-scanner (Cytiva) and quantified by MultiGauge V3.2 software (Fujifilm). The resulting electrophoretic mobility profile was fitted using OriginPro 2023.

### Supershift EMSA

Proteins were diluted in Dilution Buffer supplemented with 2 mM MgCl_2_ and 2 mM ATP. RAD51 (0.6 µM) was incubated with 5′ FITC labelled 61mer oligonucleotide (0.6 µM, concentration in nucleotides) in the presence of 10 mM Tris, pH 7.5, 10 mM KCl, 1 mM DTT, 2 mM MgCl_2_, 2 mM ATP for 5 min at 37°C. CX3 (final concentration 0.6 µM, diluted in Dilution Buffer supplemented with 2 mM MgCl_2_ and 2 mM ATP) was added followed by further 5 min incubation. Binding was terminated by the addition of 0.5 µL of 0.5 % glutaraldehyde for 10 min. Crosslinking was quenched by the addition of 1 µL of 1 M Tris, pH 8.0. Crosslinked protein-DNA complexes were incubated with 0.3 µL of anti-XRCC3 antibody (Abcam, ab20254) for 10 min at 37°C and products were resolved using 0.8% TBE agarose gel at 4°C for 60 min (6.5 V/cm).

### DNAse I protection assay

Proteins were diluted from concentrated stocks into Dilution Buffer supplemented with 2 mM MgCl_2_ and 2 mM ATP (or indicated nucleotide cofactor) which was also used in no protein controls. RAD51 (or Walker box variants, 0.23 μM or 0.6 μM each) was incubated with 30 nM 5’-FITC-labelled 61mer oligonucleotide in 10 mM Tris-HCl (pH 7.5), 10 mM KCl, 1 mM DTT, 2 mM ATP (or indicated nucleotide cofactor), 2 mM MgCl_2_, 1 mM CaCl_2_, in 10 μL reaction volume at 37°C for 5 min. Then, CX3 in indicated concentrations was added followed by further 5 min incubation. Bovine pancreatic DNaseI (1 μL (2 U), New England Biolabs) was added for 20 min at 37°C. The samples were deproteinized with 0.125% SDS and 15.6 μg proteinase K for 10 min at 37°C and resolved in 10% native polyacrylamide gels in 1X TBE (110 V, 50 min) SE206 (Hoefer) apparatus. Gels were imaged on a FLA-9000 scanner (Fujifilm) and quantified with Multi Gauge V3.2 (Fujifilm).

### ATPase assay

RAD51 or its Walker box variants, CX3 or both proteins at the indicated concentrations were mixed with A-Buffer (50 mM Tris, pH 7.5, 50 mM KCl, 2 mM MgCl_2_) in the presence or absence of (dT)63 oligonucleotide, respectively. Mixtures were pre-incubated on ice for 15 minutes. Reactions were started by the addition of ATP up to 100 µM in the final volume followed by incubation at 37°C. Each time point, 50 µL of reaction was withdrawn and mixed with 12.5 µL of Goldmix/Accelerator (1:100) (Innova Biosciences PiColorLock Gold Colorimetric Assay kit) followed by 5 min incubation at 25°C. Then, 5 µL of Stabilizer was added for further 30 min. Standard curve was constructed according to manufacturer’s instructions. Fluorescence signal was measured using Infinity F500 microplate reader (Tecan Group Ltd.) in 96-well plates at 25°C.

### Capture assay

Proteins were diluted in Dilution Buffer supplemented with 2 mM MgCl_2_/2 mM ATP (or indicated nucleotide cofactor). RAD51 was incubated with CX3 for 15 min on ice prior to the addition of Capture Buffer (35 mM Tris-HCl (pH 7.5), 50 mM KCl, 1 mM DTT, 2 mM ATP or indicated nucleotide cofactor, 2 mM MgCl_2_) and 5 µL of streptavidin coated magnetic beads (Dynabeads M-280 Streptavidin, Invitrogen) bound to either 5′ or 3′-biotinylated (dT)43 ssDNA (3′ or 5′-FITC labelled respectively). DNA conjugation to the beads was performed according to manufactureŕs instructions in 10 µL final reaction volume. Proteins were bound for 10 min at 37°C, and the beads were captured with the Magnetic Particle Separator (Roche Applied Science). Individual supernatants were set aside for further analysis. After washing the beads twice with 50 µL of Capture Buffer (2 mM ATP or indicated cofactor). Bound proteins were eluted from the beads with 10 µL of Laemmli Buffer (2% SDS, 5% 2-mercaptoethanol, Bromophenol blue). Supernatants and SDS eluates were analysed on 12% SDS-PAGE gel using SE206 (Hoefer) apparatus, followed by Coomassie Blue staining and western blotting using anti-XRCC3 (Abcam, ab20254), anti-RAD51C (Novus, NB100-177SS) and anti-RAD51 (Santa Cruz, H-92) antibodies, respectively.

### Biolayer-interferometry assay (BLI)

BLI experiments were performed using a single-channel BLItz instrument in Advanced Kinetics mode (Sartorius) at room temperature and with continuous shaking at 2.200 rpm. Prior to the measurements, streptavidin biosensors (SAX, Sartorius) were pre–hydrated by incubation with SF buffer with 0.05% Tween 20 for 10 min. Subsequently, the biosensor was loaded with 15 nM biotinylated dT ssDNA for 120 seconds and washed with SF buffer. Next, a solution containing RAD51, CX3 or mixture of both proteins was applied to the biosensor in SF buffer containing 1 mM ATP and 0.05% Tween-20, and association was monitored for 180 seconds. The binding affinity of the proteins to DNA was determined by measuring the increase in the thickness of the biomolecule layer in nanometers (nm). To monitor the dissociation of protein.ssDNA filaments, 10 µM BRC4 peptide in SF buffer was mixed with filaments in dissociation phase. For CX3 and RAD51 protein interaction experiments Ni-NTA biosensors (Sartorius) were used, SF buffer containing 1 mM ATP, 0.05% Tween-20 and 15 mM imidazole was used for loading, association and dissociation steps.

### Pulldown assay

To detect the interaction between CX3 and RAD51, we mixed 2.5 µg of purified CX3 (containing HIS-tagged RAD51C) and 2.5 µg of purified RAD51 together with 7.5 µL of Ni-NTA beads. The reaction was prepared in total volume of 20 µL of SF buffer supplemented with 1 mM ATP (AMP-PNP, ADP), 15 mM imidazole and 0.05% Tween-20, and incubated on ice for 1 hr. Subsequently, the beads were then washed 2 times with the same buffer, to remove any non-specifically bound proteins. Bound proteins were eluted by boiling beads in Laemmli Sample Buffer.

### Microscale thermophoresis

Binding affinity quantifications via microscale thermophoresis were performed using the Monolith NT.115 (Nanotemper Technologies). Affinity measurements were performed in SF buffer containing 0.05% Tween-20 plus 1 mM ATP. Samples were loaded into Premium capillaries (NanoTemper Technologies). Measurements were performed at 25°C, 40 % Excitation power, medium MST-power and constant concentration of 40 nM fluorescently labelled RAD51 protein (Protein Labeling Kit RED-maleimide 2nd Generation, NanoTemper technologies) and increasing concentration of purified CX3 protein. Data were analysed by the MO. Affinity Analysis software (NanoTemper Technologies).

### Stopped-flow measurement

Stopped-flow experiments were performed similarly as described previously^33^ using an SFM-300 stopped-flow machine (Bio-Logic) fitted with a MOS-200 monochromator spectrometer (Bio-Logic) with excitation wavelength set at 545 nm at 37°C. Fluorescence measurements were collected with a 550 nm long pass emission filter. All reactions were performed in Stopped Flow Buffer (50 mM Tris-HCl (pH 7.5), 10 mM MgCl_2_, 50 mM NaCl, 1 mM ATP) with 15 nM 5’-Cy3 fluorescently labelled (dT)43 (Cy3-43mer). In experiments with pre-incubation step, proteins or proteins with DNA were incubated for 5 minutes at 37°C before stopped-flow set-up. Proteins were added directly from concentrated stocks. Unlabelled (dT)43 or (dT)79 was used at 1500 nM (moles) (100-fold excess) in competition experiments. For all experiments, controls were also performed for buffer alone with and without DNA to confirm fluorescence signal stability over the time course of the experiments (data not shown). We verified that the theoretical minimum concentration of RAD51 (350 nM) required to fully coat all DNA molecules (645 nM nucleotides in stopped-flow) with filaments was sufficient to induce a change in Cy3 fluorescence over longer time courses. Fluorescence measurements for most experiments were collected according to the following protocol: (1) every 0.00005 s from 0-0.05 s; (2) every 0.0005 s from 0.05-0.56 s; (3) every 0.02 s from 0.56-60.54 s.

### Electron microscopy

To measure length of RAD51-ssDNA filament, RAD51 (4 µM) was pre-incubated with indicated concentration of CX3 (from 0.04 to 1 µM) for 1 min at RT. Then RAD51 or RAD51-CX3 was incubated with 133 nM Cy3-labelled 90mer oligonucleotide (12 µM in nucleotides, Sigma Aldrich or Merck) in 50 mM Tris-HCl pH 7.5, 10 mM KCl, 10 mM MgCl_2_ and 1 mM ATP for 10 min at 37°C. Reaction was finally cross-linked with 0.01% glutaraldehyde. Grids were discharged for 20 s (hydrogen-oxygen) in Gatan Solarus II. Sample (4 µl) of the reaction was placed on discharged grids, incubated for 1 min followed by staining with 2% uranyl acetate for 1 min. Micrographs were collected with transmission electron microscope Talos F200C (details). The length of RAD51-(CX3)-ssDNA filaments was analysed blindly using ImageJ. For each condition, at least 150 filaments were analysed.

The RAD51-ssDNA complexes were formed in buffer containing 50 mM Tris-HCl pH 7.5, 50 mM NaCl, 10 mM MgCl2 and 1 mM ATP. RAD51 (5.4 µM) alone or with CX3 (1.1 µM) protein was mixed with 16.2 µM ssDNA 150mer (concentration in nucleotides) and incubated for 10 min at 37°C. The ssDNA 150mer oligonucleotide was purchased from VBC Biotech. A sample of the mixture (4 µL) was deposited on glow-discharged grids coated with 12 nm layer of homemade continuous carbon and stained with Nano-W (Nanoprobes Inc.). Data were collected on transmission electron microscope Tecnai F20 (FEI, Eindhoven) operating at 200 keV with a nominal magnification of 50,000x resulting in pixel size of 2.22 Å/px. The data were acquired on CCD camera (FEI Eagle) under low-dose conditions (∼25 e-/Å2s) and with the under-focus in the range of 2.5 – 4.5 µm. Immuno-gold labelling experiments were performed as described previously^34^. Anti-XRCC3 10F1 antibody (Abcam, ab20254) was conjugated to 20-nm gold particles using the InnovaCoat Gold Conjugation Mini kit (Innova Biosciences). For quantification, discrete filaments were counted and scored for number bound with anti-XRCC3-20-nm gold on the end or along the filament length.

### Quantification and Statistical Analysis

For EMSA DNA binding experiments, FITC-labelled DNA fluorescence was quantified with Multi Gauge V3.2 (Fujifilm). After adjusting for background, the % of DNA binding was assessed as a ration between fluorescence of free DNA signal to no-protein signal. Average relative binding values from at least 3 independent experiments were then determined. Error bars represent standard deviation. For nuclease protection experiments, after adjusting for background, the % protection was determined for all samples then normalized to the value for “minus DNaseI control“. Average relative protection values from at least 3 experiments were then determined. Error bars represent standard deviation. For ATPase activity, values reported represent averaged data from at least 3 independent experiments, error bars represent standard deviation. In stopped-flow experiments, traces were collected from between three and six independent reactions (n = 3-6, see individual figure legends for details). For presentation, average traces for each experiment were generated. Importantly, all conclusions from this study are made based on the magnitude of changes in fluorescence over time and how these vary with CX3 concentration, and therefore the absolute fluorescence values were converted to arbitrary units by a normalization procedure to facilitate comparison. For all experiments the raw data were normalized to the same fluorescence value at the 0.01 s time point since an initial fluctuation in fluorescence was observed in most samples, to decrease background noise. These adjustments were to aid comparison of protein concentration-dependent trends in the change in fluorescence over time. An exception being RAD51 filament formation experiments (Fig. S4B) where raw data were normalized for the 0 s time point. For analysis, for all experiments twenty-point moving averages were calculated on each individual normalized trace, which were used to define initial (0 s or 0.01 s) and final (60.54 s) Cy3 fluorescence values for each experiment. Average values and associated standard deviations were then calculated.

## ACKNOWLEDGEMENTS

The work in L.K. group was supported by the Czech Science Foundation (21-22593X and 25-17326S) and the European Union’s Horizon Research and Innovation Programme [101158508] TRIAD. Z.P. acknowledges support by the EU’s Horizon Europe Programme under the Grant agreement No. 101136607 (CLARA Project), and by the EU from the Operational Programme Jan Amos Komenský (project “Center for Artificial Intelligence and Quantum Computing in System Brain Research”, reg. no. CZ.02.01.01/00/23_029/0008437). Work in G.J.W. group was supported by the Canadian Institutes of Health Research (PJT159676) and National Institute of Health 7R21ES025895). Work in J.A.T. group was funded by the National Institute of Health (PO1CA92584 to G.J.W. and J.A.T.). J.A.T. efforts were also supported by R35 CA220430 and Robert A. Welch Chemistry Chair G-0010. CIISB research infrastructure project LM2015043 funded by MEYS CR is acknowledged for the financial support of the measurements at the Cryo-electron Microscopy and Tomography core facility. Access to computing and storage facilities of the National Grid Infrastructure MetaCentrum, provided under the programme CESNET LM2015042, is greatly appreciated.

## AUTHOR CONTRIBUTIONS

O.B. performed most of the *in vitro* assays and analysed the data; M.S. performed the stopped flow, BLI, SEC, pull-down experiments and analysed the data; J.M. performed the immunogold EM. O.B., G.J.W., J.P. and M.S. cloned, expressed and purified the proteins; L.M. and M.B.P. carried out some of the *in vitro* assays and analysed data; G.J.W. and J.A.T. oversaw and conceived the strategy for recombinant human CX3 production and analysis of its function; L.K. conceived of and oversaw the project; Z.P. constructed the kinetic model, performed numerical simulations, and contributed to the manuscript preparation and discussion; M.S., O.B., and L.K. wrote the paper; all authors discussed and edited the manuscript.

## CONFLICT OF INTEREST

The authors declare that they have no known competing financial interests or personal relationships that could have appeared to influence the work reported in this paper.

## Notes

### Competing Interest Statement

The authors have declared no competing interest.

