## Supplementary material for "Filament assembly, stabilisation, and ATP-dependent RAD51 filament capping and remodeling by RAD51C-XRCC3 complex"

### *Supplementary figures*

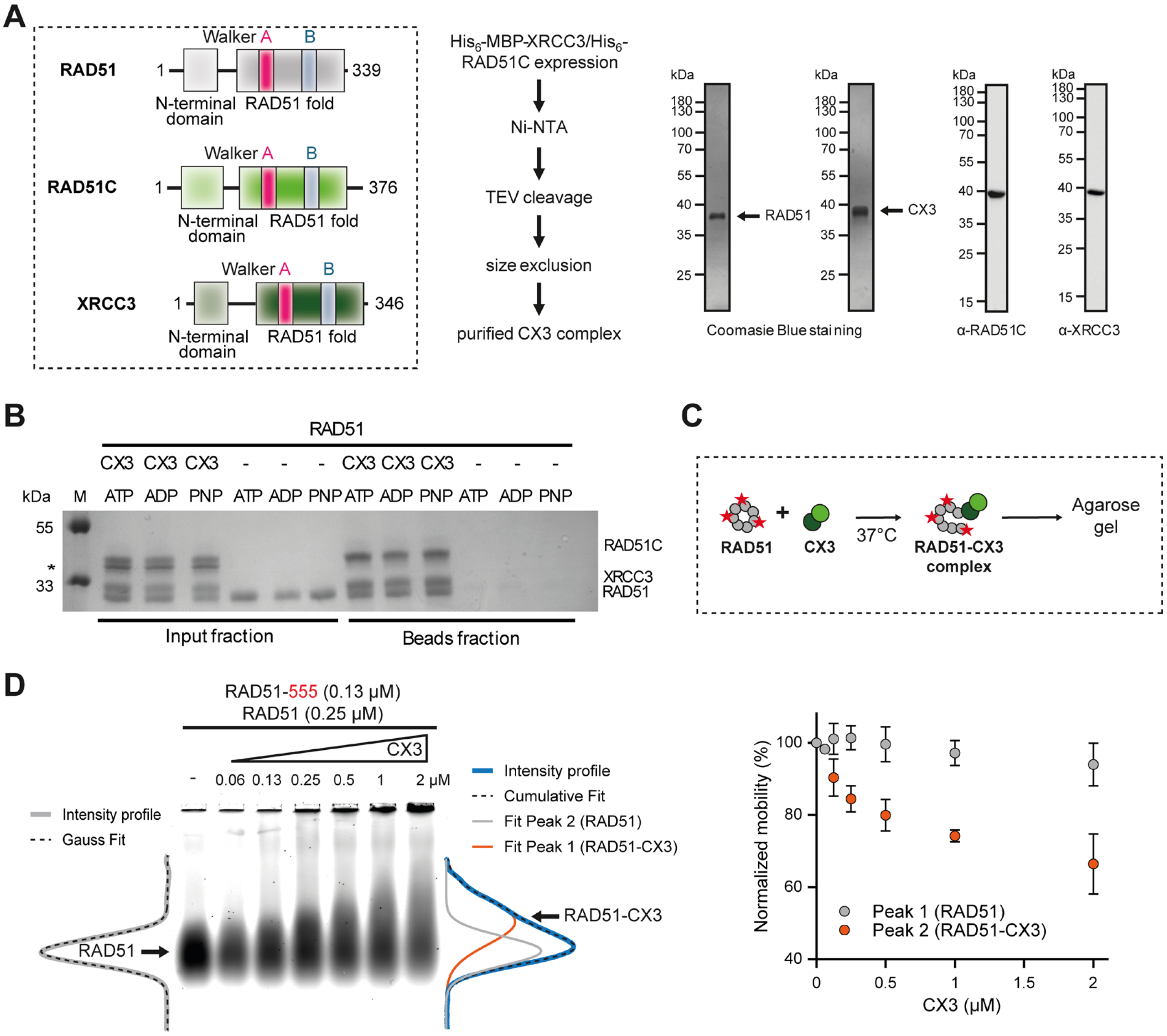


**Fig. S1: CX3 purification and its interaction with RAD51. (A)** Domain organisation of human RAD51, RAD51C and XRCC3 (left). Schematics of the purification strategy for CX3 complex (middle). Coomassie Blue-stained SDS-PAGE of purified RAD51 (0.5 μg) and CX3 complex (1.0 μg complex, ~ 0.5 μg each component) used in the study. Western blot (α-RAD51C, α-XRCC3) of purified CX3 complex (right). **(B)** Size-exclusion chromatography profile of CX3 complex during purification. **(C)** Pull down analysis of RAD51 (0.5 μg) and CX3 (0.5 μg) in the presence or absence of indicated nucleotide cofactors. **(D)** Schematics of the EMSA used to analyse CX3-RAD51 interaction. Red asterisk represents Alexa Fluor 555 dye. **(E)** Representative image of the EMSA analysis (left) together its quantification (right). Error bars represent mean ±

s.d. (n = 3). Lines beside the gels show the fluorescence intensity profiles of RAD51 alone (grey) and in the presence of CX3 (2 μM, blue). Dashed lines indicate the fits of the profiles.

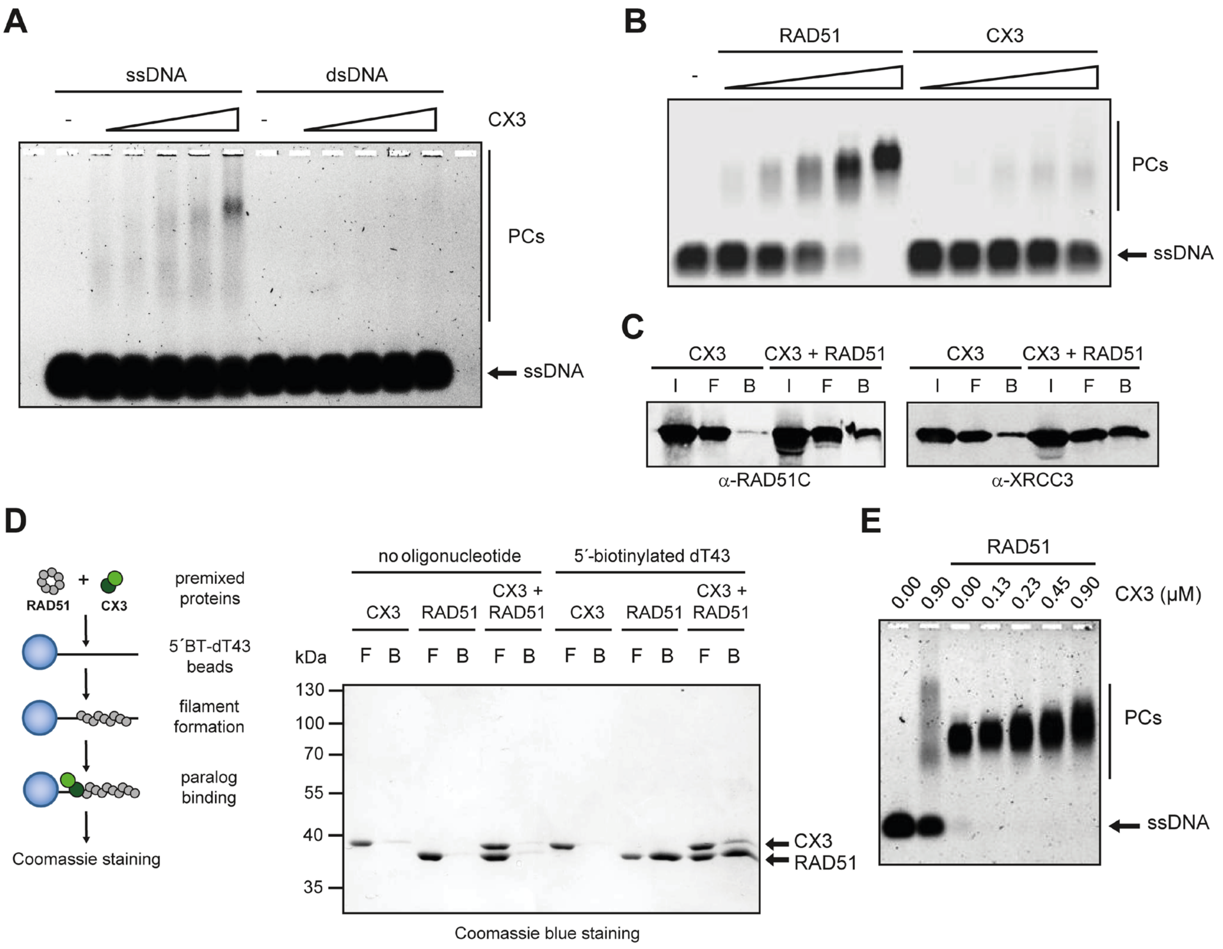


**Fig. S2: Additional characterisation of CX3 DNA binding and association with RAD51 filaments. (A)** Representative EMSA binding of CX3 (0.09-1.470 μM) to 30 nM 49mer ssDNA or dsDNA. Protein-DNA complexes (PCs) were stabilised by crosslinking with 0.5 μL of 0.2% glutaraldehyde and resolved on 0.8% TBE agarose gel. (**B**) Representative EMSA comparing the binding of RAD51 (0.056-0.9 μM) and CX3 (0.056-0.9 μM) to 30 nM 61mer ssDNA. Formed complexes were stabilised by crosslinking with 0.5 μL of 0.07% glutaraldehyde and analysed on 0.8% TBE agarose gel. (**C**) Representative Western blot analysis of the capture assay shown in Fig. 1E using α-RAD51C or α-XRCC3 antibodies. Protein input (I), flow-through (F) and bound (B) fractions are indicated. (**D**) Schematics of the capture assay (left) and representative SDS-PAGE gel (right) with purified RAD51 (6.5 μM) and/or CX3 (0.94 μM) using 5´ biotinylated (BT) ssDNA or DNA-free streptavidin coated beads as a control. Proteins were incubated with magnetic beads for 10 min at 37°C. Flow-through (F) fraction was collected followed by two washing steps and protein elution from the beads (B). (**E**) Representative EMSA of RAD51 (0.90 μM) assembled on 30 nM 61mer ssDNA in the absence or presence of increasing CX3 concentration (0.13-0.9 μM).

Complexes were crosslinked with 0.5 µL of 0.2% glutaraldehyde and resolved on 0.8% TBE agarose gel.

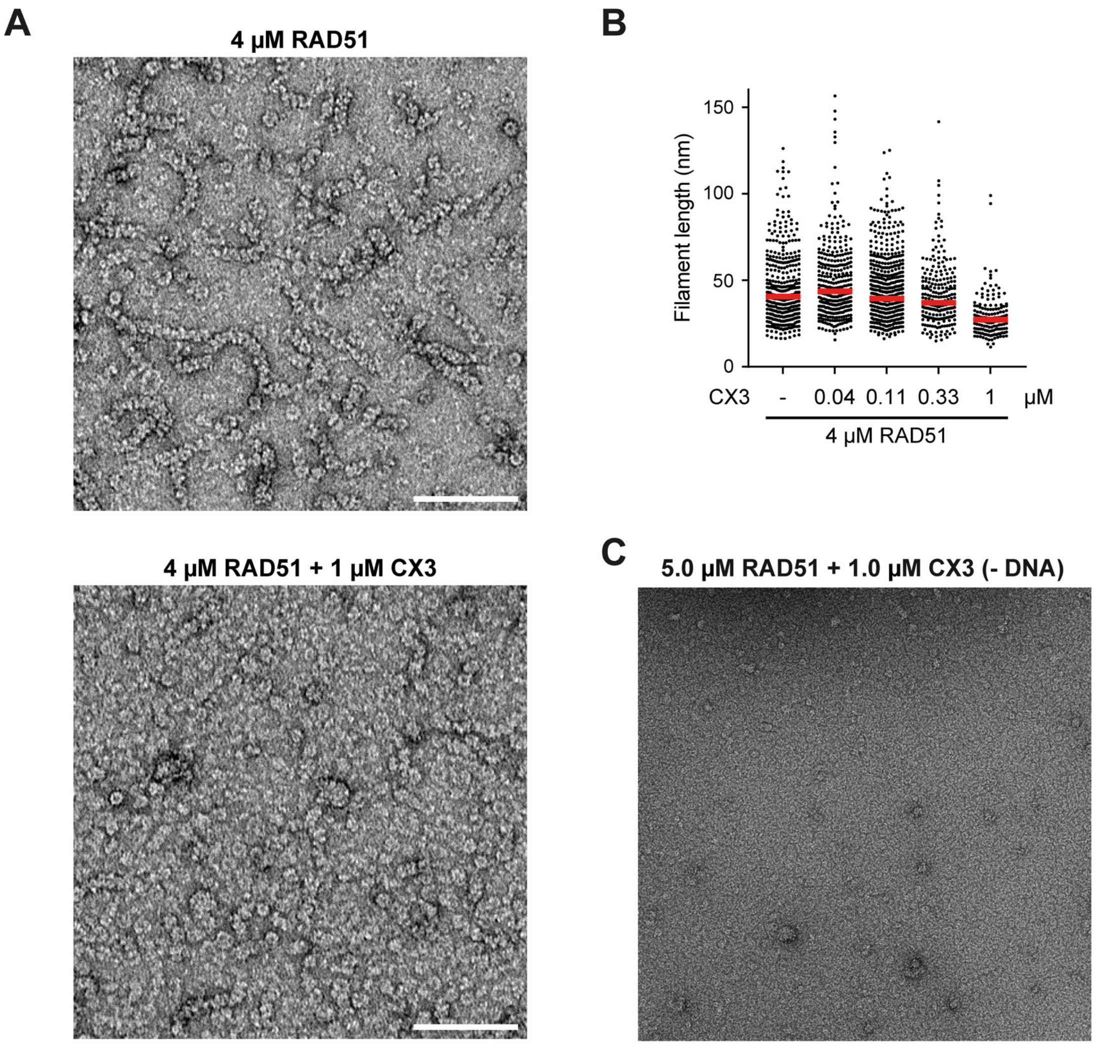


**Fig. S3: Negative-stain electron microscopy (EM) analysis.** (A) Representative negative-stain EM of RAD51 (4 µM) filaments formed on 90mer ssDNA in the absence (top) or presence (bottom) of CX3 (1 µM). Scale bar, 100 nm. (B) Quantification of the length of RAD51 filaments formed on 90mer ssDNA in the presence or absence of CX3 (0.04-1 µM), analysed by negative-stain EM. At least 150 RAD51 filaments were evaluated for each condition. (C) Representative negative-stain EM of RAD51 and CX3 in the absence of DNA.

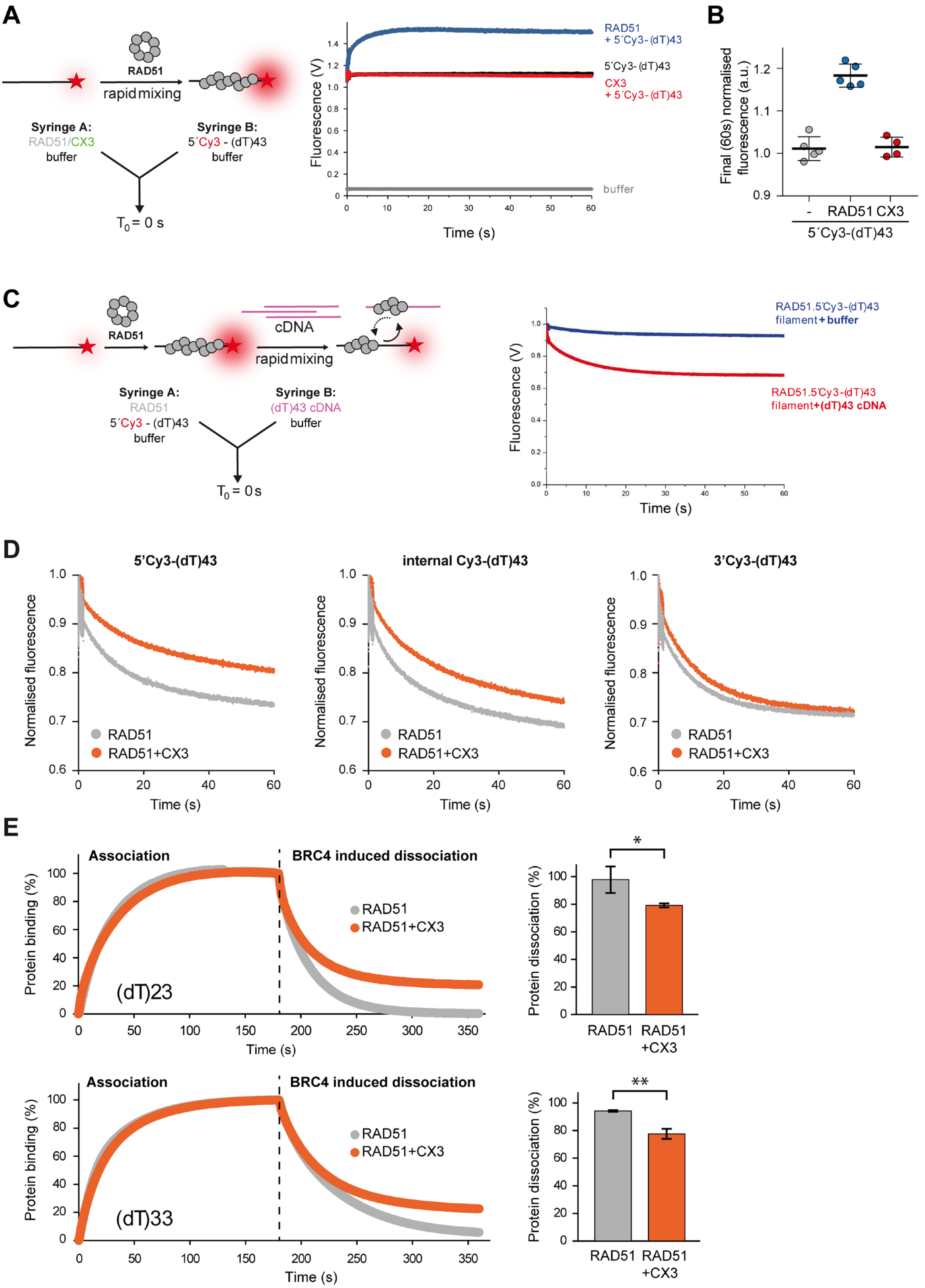


**Fig. S4: Additional characterisation of CX3-mediated RAD51 filament stabilisation. (A)** Schematic of stopped-flow filament assembly experiment (left).

RAD51 (0.350 μM) or CX3 (0.175 μM) was pre-incubated with 1 mM ATP for 5 min and rapidly mixed with 15 nM 5´-Cy3 labelled (dT)43 ssDNA. Representative traces are shown for buffer (grey), Cy3-labelled ssDNA (black), and RAD51 (blue) and CX3 (red) binding to DNA (right). Average normalised fluorescence was plotted as a function of time. **(B)** Quantification of normalised endpoint (60s) Cy3 fluorescence for the data presented in (A). Error bars indicate ± s.d. (n = 4-5). **(C)** Schematic of stopped-flow filament disassembly experiment (left). RAD51 (0.350 μM) was pre-incubated with 1 mM ATP for 5 min, followed by incubation with 15 nM 5´-Cy3 labelled (dT)43 ssDNA for further 5 min to assemble RAD51-ssDNA filaments. Pre-formed RAD51 filaments were rapidly mixed with 100-fold excess (1.5 μM) of unlabelled competitor (dT)43 ssDNA (cDNA). Dissociation of RAD51 from ssDNA results in a decrease in Cy3 fluorescence. Representative traces are shown for RAD51 filaments mixed with buffer (blue) or competitor DNA (red) (right). Average normalised fluorescence was plotted as a function of time. **(D)** Average stopped-flow fluorescence intensity traces monitoring the stability of RAD51 filaments assembled on (dT)43 ssDNA labelled with Cy3 at the 5´ end (left), internally (middle), or at the 3' end (right). RAD51 (1 μM), with (orange) or without (grey) CX3 (0.334 μM), was pre-incubated with 15 nM labelled DNA for 10 min and then mixed with a 100-fold excess (1.5 μM) of unlabelled (dT)43 competitor ssDNA. Data represent (n = 3-4) (**E**) Representative bio-layer interferometry (BLI) sensorgrams (left) of RAD51 filament stability on short (dT)23 and (dT)33 ssDNA substrates. RAD51 (0.75 μM) was bound to streptavidin biosensors carrying 15 nM 3′-biotinylated (dT)23 (top) or (dT)33 (bottom) ssDNA in the absence (grey) or presence of CX3 (0.75 μM, orange). Dissociation was facilitated by addition of 10 μM BRC4 peptide. Endpoint normalised optical thickness is shown (right). Error bars indicate ± s.d. (n = 3). Statistical significance was determined using two-tailed Student's t test: * $p < 0.05$, ** $p < 0.01$.

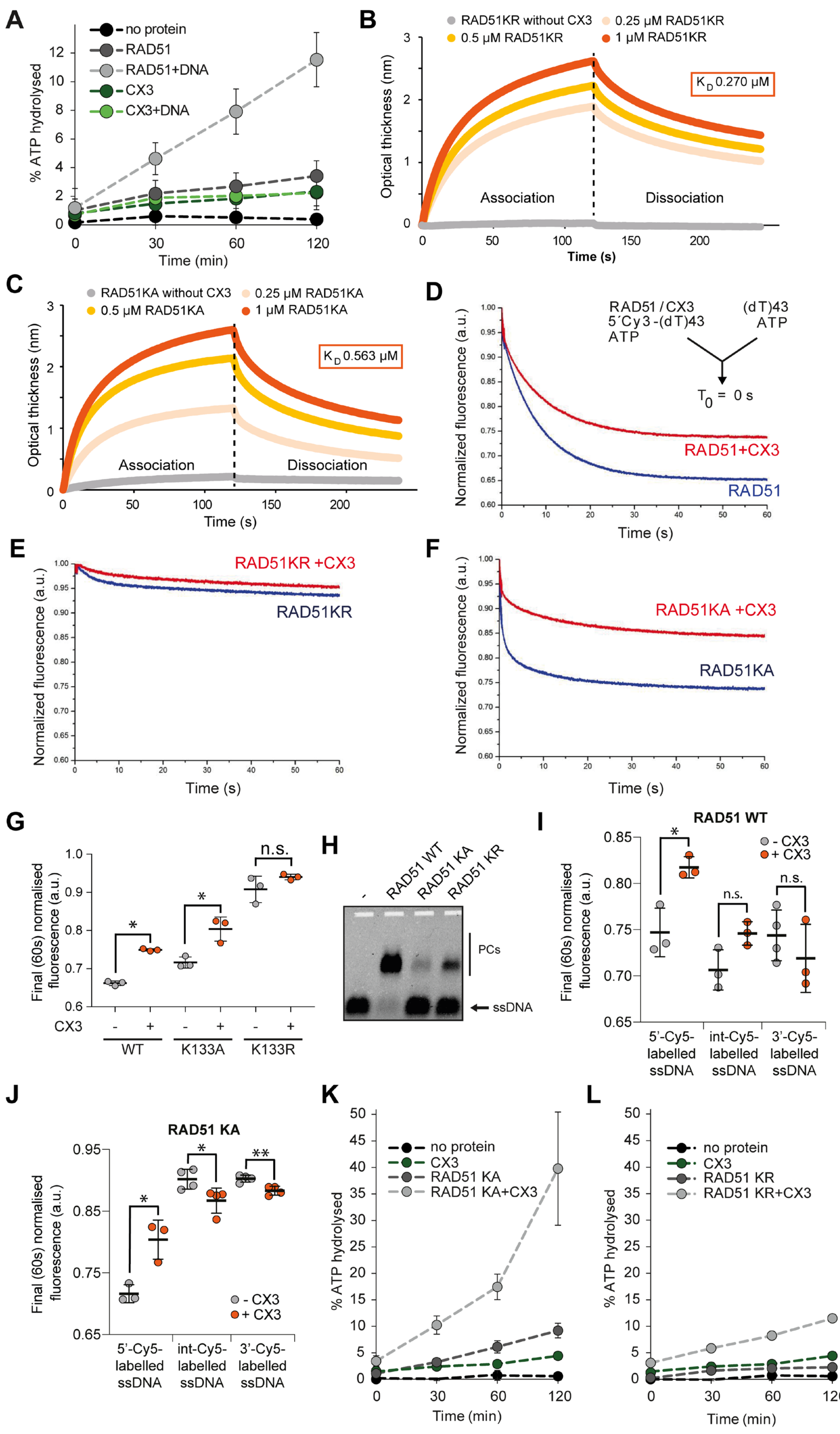
A
% ATP hydrolysed
Time (min)
no protein
RAD51
RAD51+DNA
CX3
CX3+DNA
B
RAD51KR without CX3
0.25 µM RAD51KR
0.5 µM RAD51KR
1 µM RAD51KR
Optical thickness (nm)
K D 0.270 µM
Association
Dissociation
Time (s)
C
RAD51KA without CX3
0.25 µM RAD51KA
0.5 µM RAD51KA
1 µM RAD51KA
K D 0.563 µM
D
RAD51 / CX3
5´Cy3-(dT)43
ATP
(dT)43
ATP
T0 = 0 s
Normalized fluorescence (a.u.)
RAD51+CX3
RAD51
E
RAD51KR +CX3
RAD51KR
F
RAD51KA +CX3
RAD51KA
G
Final (60s) normalised fluorescence (a.u.)
n.s.
CX3
WT
K133A
K133R
H
RAD51 WT
RAD51 KA
RAD51 KR
PCs
ssDNA
I
RAD51 WT
- CX3
+ CX3
5'-Cy5-labelled ssDNA
int-Cy5-labelled ssDNA
3'-Cy5-labelled ssDNA
J
RAD51 KA
K
no protein
CX3
RAD51 KA
RAD51 KA+CX3
L
no protein
CX3
RAD51 KR
RAD51 KR+CX3

**Fig. S5: Additional characterisation of the effect of CX3 on RAD51 Walker mutants. (A)** ATPase activity of RAD51 (0.250 µM) or CX3 (0.250 µM) measured in the presence (light grey or green) or absence (dark grey or green) of 2.5 µM (dT)63 ssDNA. Error bars indicate ± s.d. (n = 3). (**B, C**) Representative BLI sensorgrams of interaction between CX3 and RAD51 Walker A mutants. His-tagged CX3 (500 nM) was immobilised on NTA-biosensors and RAD51 K133R (KR) (**B**) or K133A (KA) (**C**) was loaded onto the biosensor. Association and dissociation phases were monitored in real time. Endpoint normalised optical thickness and calculated $K_D$ are shown. **(D-F)** Stopped-flow analysis of RAD51 filament stability. RAD51 WT (**D**), KR (**E**) or KA (**F**) (0.700 µM each) was pre-incubated with 15 nM 5´Cy3-(dT)43 with (red) or without (blue) CX3 (0.175 µM) for 10 min before rapid mixing with 100-fold excess (1.5 µM) of unlabelled (dT)43 competitor ssDNA. Average normalised Cy3 fluorescence is plotted as a function of time. (**G**) Quantification of the normalised endpoint (60s) Cy3 fluorescence for the data presented in D-F. Error bars indicate ± s.d. (n = 3). Statistical significance was determined using two-tailed Student's t test: n.s. not significant, * $p < 0.05$. **(H)** Representative EMSA binding of RAD51 WT, K133A or K133R (0.23 µM) to 30 nM 61mer ssDNA. Protein-DNA complexes were stabilised by crosslinking with 0.5 µL of 0.01% glutaraldehyde and resolved on 0.8% TBE agarose gel. **(I)** RAD51 (1 µM) or its KA **(J)** variant with (red) or without (blue) CX3 (334 nM) was pre-incubated with 15 nM 5´Cy3-(dT)43, (dT)10-Cy3(dT)-(dT)32 or 3´Cy3-(dT)43 for 10 min and then mixed with excess (1.5 µM) of unlabelled (dT)43 ssDNA. Quantification of normalized end point (60 s) Cy3 fluorescence for different conditions. Error bars indicate s.d. (n = 3 - 4). Statistical significance was determined using two-tailed Student's t test: n.s. not significant, * $p < 0.05$. ** $p < 0.01$. **(K, L)** ATPase activity of CX3 (0.400 µM) in the presence of ATPase-deficient RAD51 Walker A box mutants. CX3 ATP hydrolysis was measured after incubation with RAD51 K133A (**K**) or K133R (**L**) (1.0 µM each) and 160 nM (dT)63 ssDNA. Error bars indicate ± s.d. (n = 4).

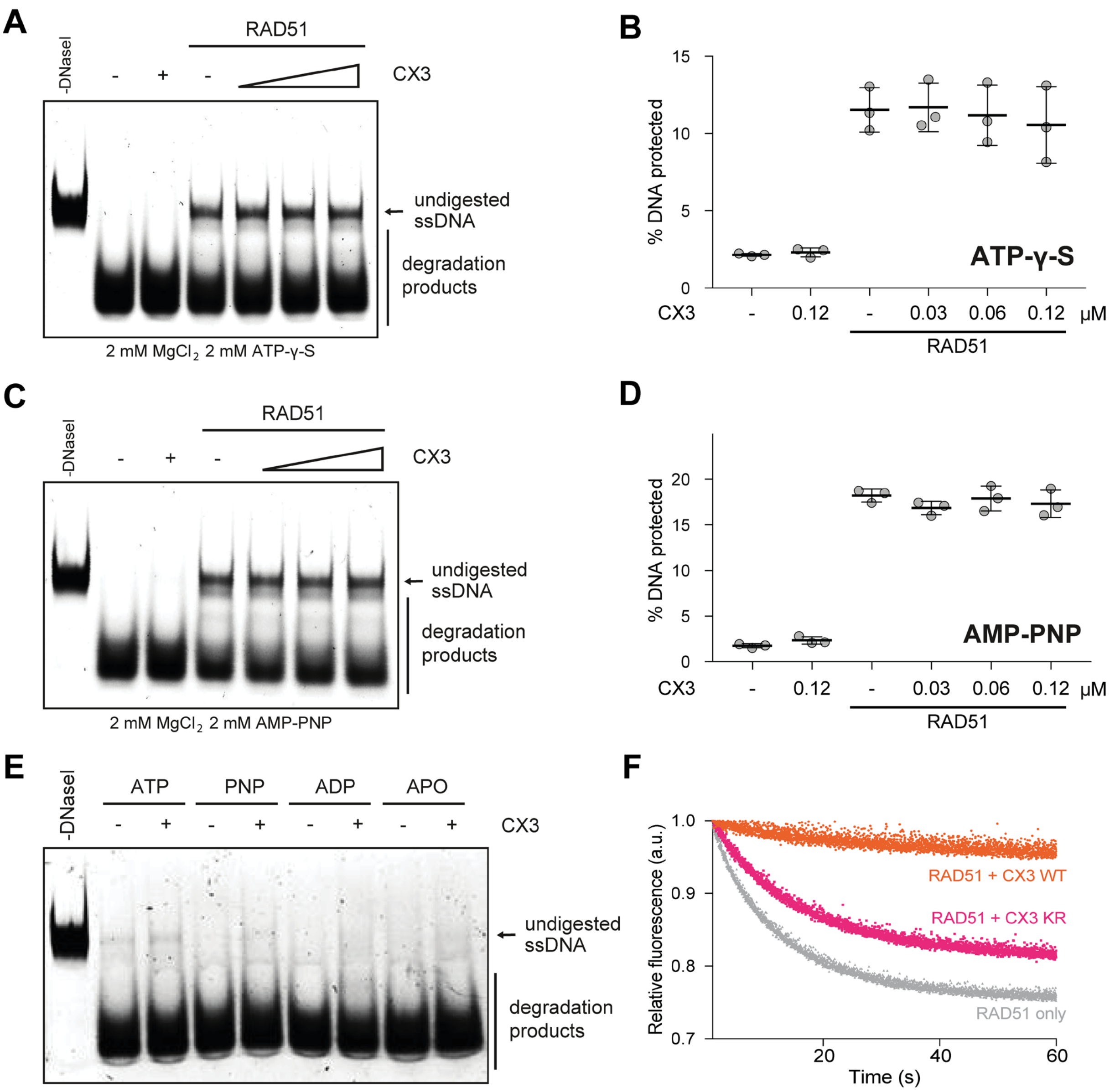


**Fig. S6: Additional characterisation of the role of ATP hydrolysis in RAD51 filament remodeling. (A)** DNase I protection assay performed in the presence of ATP-γ-S for RAD51 (0.12 μM) and/or CX3 (0.03 - 0.12 μM range). Proteins were incubated with 30 nM 61mer ssDNA for 10 min at 37°C. Protein-DNA complexes were challenged with 1 U of DNase I for 20 min at 37°C. Reactions were terminated using 0.125% SDS and 15.6 μg proteinase K and resolved on 10% polyacrylamide gel. **(B)** Quantification of data from panel (A). Error bars indicate ± s.d. (n = 3). **(C)** DNase I protection assay performed similarly to (A) in the presence of AMP-PNP. **(D)** Quantification of data from panel (C). Error bars indicate ± s.d. (n = 3). **(E)** DNase I protection assay of CX3 alone performed similarly to Fig. 5B. CX3 (0.23 μM) was incubated with 30 nM 61mer ssDNA in the presence of indicated nucleotide cofactors for 10 min at 37°C before challenge with 1 U of DNase I for 20 min at 37°C. Reactions were terminated using 0.125% SDS and 15.6 μg proteinase K and resolved on 0.8% TBE agarose gel. **(F)** Stopped flow analysis of RAD51 stability in the presence of CX3 WT or CX3 KR. RAD51 (500 nM) was pre-incubated with 15 nM 5´-Cy3-(dT)43 ssDNA with or without CX3 WT or CX3 KR (250 nM) for 10 min before rapid mixing with 100-fold excess of unlabelled (dT)43 ssDNA. Average normalised Cy3 fluorescence traces are shown.

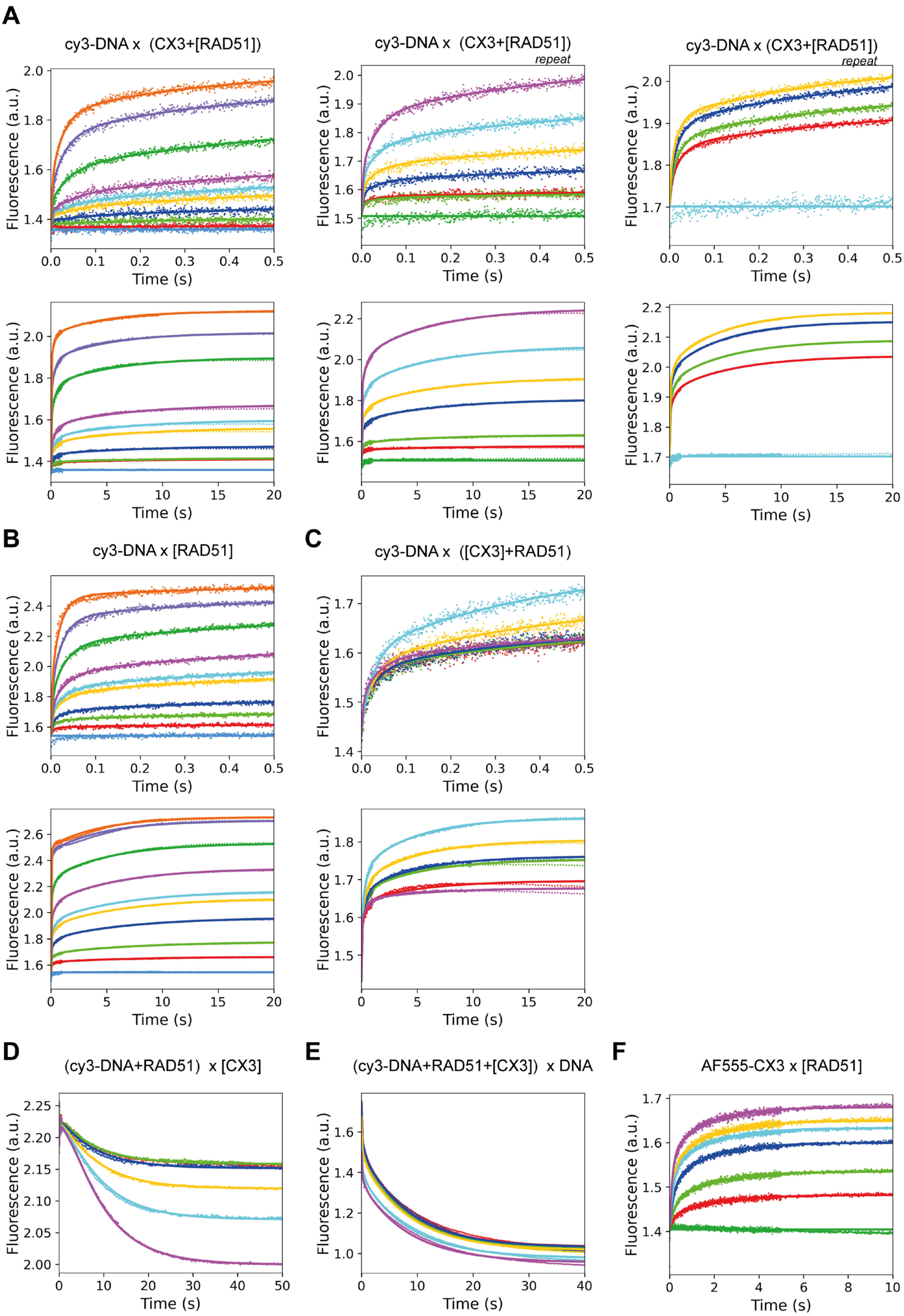


**Fig. S7: Experimental traces from global kinetic binding analysis.** Stopped flow fluorescence traces (excitation 545 nm, emission > 550 nm) recorded upon mixing reaction components depicted above each chart. **(A)** RAD51 titration at a fixed CX3 concentration followed by addition of Cy3-labelled (dT)43 ssDNA. **(B)** RAD51 titration with Cy3-labelled dT43 ssDNA in the absence of CX3. **(C)** CX3 titration at a fixed RAD51 concentration followed by addition of Cy3-labelled (dT)43 ssDNA. **(D)** CX3 titration to pre-assembled RAD51-Cy3-labelled (dT)43 ssDNA filaments. **(E)** Competitor-DNA challenge of pre-formed RAD51-Cy3-labelled (dT)43 ssDNA filaments in the presence and absence of CX3. **(F)** RAD51 titration with fluorescently-labelled CX3.

Each trace represents an average of 3 to 6 measurements. The solid black lines represent the best global fit to the kinetic model to the experimental data. The total fluorescence signal was calculated as the sum of the contributions from all fluorescent species present throughout the reaction. The contribution of each species was assumed to be proportional to its concentration and was described by scaling functions (Equations 1–3). The corresponding scaling factors were globally optimized and shared across all experiments.

**A**

| Parameter | Unit | RAD51-CX3<br>**Value ± s.e.**<br>lower; upper limit |
|---|---|---|
| $k_1$ | ($nM^{-1}.s^{-1}$) | **0.060 ± 0.001**<br>0.049; 0.072 |
| $k_{-1}$ | ($s^{-1}$) | **180 ± 2**<br>144; 226 |
| $k_2$ | ($nM^{-1}.s^{-1}$) | **0.045 ± 0.001**<br>0.030; 0.108 |
| $k_{-2}$ | ($s^{-1}$) | **25.4 ± 0.3**<br>16.2; 77.4 |
| $k_3$ | ($s^{-1}$) | **3.65 ± 0.08**<br>2.33; 6.90 |
| $k_{-3}$ | ($s^{-1}$) | **2.09 ± 0.03**<br>1.39; 2.95 |
| $k_4$ | ($s^{-1}$) | **0.219 ± 0.004**<br>0.110; 0.337 |
| $k_{-4}$ | ($s^{-1}$) | **0.127 ± 0.001**<br>0.120; 0.134 |
| $k_5$ | ($nM^{-1}.s^{-1}$) | **0.0050 ± 0.0002**<br>0.0031; 0.0075 |
| $k_{-5}$ | ($s^{-1}$) | **2.93 ± 0.04**<br>2.07; 4.60 |
| $k_6$ | ($s^{-1}$) | **0.185 ± 0.009**<br>0.077; 0.340 |
| $k_{-6}$ | ($s^{-1}$) | **0.489 ± 0.005**<br>0.378; 0.616 |
| $k_7$ | ($nM^{-1}.s^{-1}$) | **0.72 ± 0.04**<br>0.38; 2.25 |
| $k_{-7}$ | ($s^{-1}$) | **50 ± 2**<br>32; 238 |
| $k_8$ | ($nM^{-1}.s^{-1}$) | **0.20 ± 0.03**<br>0.06; 0.95 |
| $k_{-8}$ | ($s^{-1}$) | **17.0 ± 0.6**<br>4.5; 41.4 |
| $k_9$ | ($s^{-1}$) | **1.5 ± 0.3**<br>0.6; 2.2 |
| $k_{-9}$ | ($s^{-1}$) | **0.5 ± 0.1**<br>0.2; 2.9 |
| $k_{10}$ | ($s^{-1}$) | **0.41 ± 0.04**<br>0.03; 1.61 |
| $k_{-10}$ | ($s^{-1}$) | **0.29 ± 0.05**<br>0.10; 2.11 |

**B**

| | RAD51 x ssDNA | RAD51+CX3 x ssDNA | Relative change |
|---|---|---|---|
| $K_{D1(7)} = k_{-1(-7)}/k_{1(7)}$ (μM) | 3.000 | 0.069 | 43 |
| $K_{D2(8)} = k_{-2(-8)}/k_{2(8)}$ (μM) | 0.564 | 0.085 | 7 |

| | RAD51 x CX3 |
|---|---|
| $K_{D5} = k_{-5}/k_5$ (μM) | 0.586 |
| $K_{D,app}$ (μM) | 0.425 |

$$K_{D,app} = \frac{k_{-5} \times k_{-6}}{k_5 \times (k_6 + k_{-6})}$$

**C**

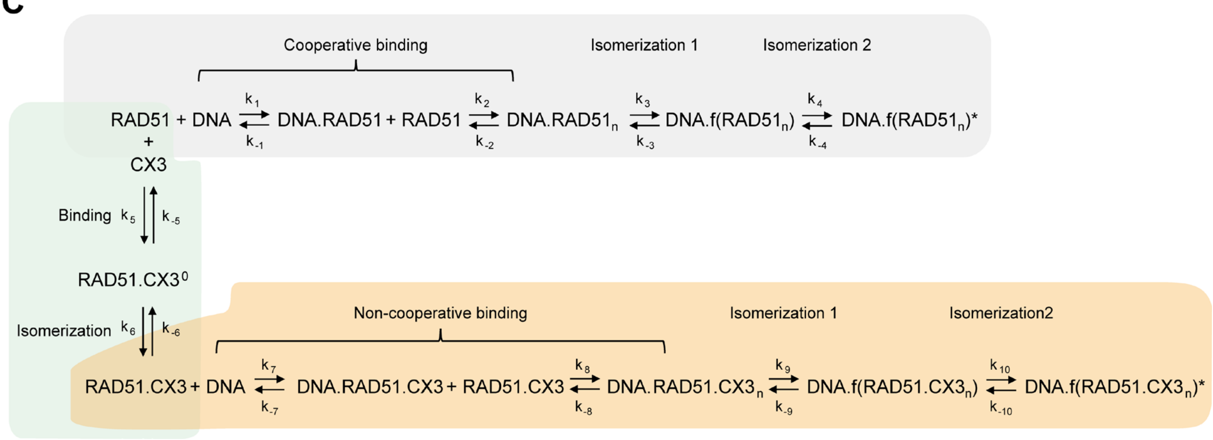


**Fig. S8: Global numerical analysis of the stopped flow kinetic data. (A)** Kinetic rate constants for individual steps determined by simultaneous fitting of the kinetic data from Fig. S7. Different scaling factors *f* were used for individual data sets reflecting variations in detector sensitivity. Defining relative change in fluorescence of individual reaction species, were determined globally and applied to all experiments. The standard error (± s.e.) was calculated from the covariance matrix during nonlinear regression. All experiments were performed in SF buffer at 37°C. **(B)** Calculated dissociation constants for protein associations with ssDNA, or association of CX3 with RAD51 using rate constants obtained from global fitting model shown in (A). **(C)** The globally fitted kinetic model summarising the interaction of RAD51 and CX3 as well as filament assembly. The individual rate constants (k) derived from the global analysis from panel (A) are indicated for each step. Orange: In the absence of CX3, RAD51 engages ssDNA through a four-step mechanism comprising cooperative binding composed of two binding events ($k_1$, $k_{-1}$) and ($k_2$, $k_{-2}$), isomerisation to the filament state ($k_3$, $k_{-3}$) and subsequent filament rearrangement ($k_4$, $k_{-4}$). Green: In the presence of CX3, RAD51 and CX3 first bind in solution ($k_5$; $k_{-5}$) to form a rearranged RAD51-CX3 complex ($k_6$; $k_{-6}$). Grey: RAD51-CX3 complex subsequently engages ssDNA through an analogous but kinetically distinct pathway consisting of two non-cooperative bindings steps and two isomerizations ($k_7$-$k_{10}$).

**A**

**RAD51(7).ATP [AF3 model]**

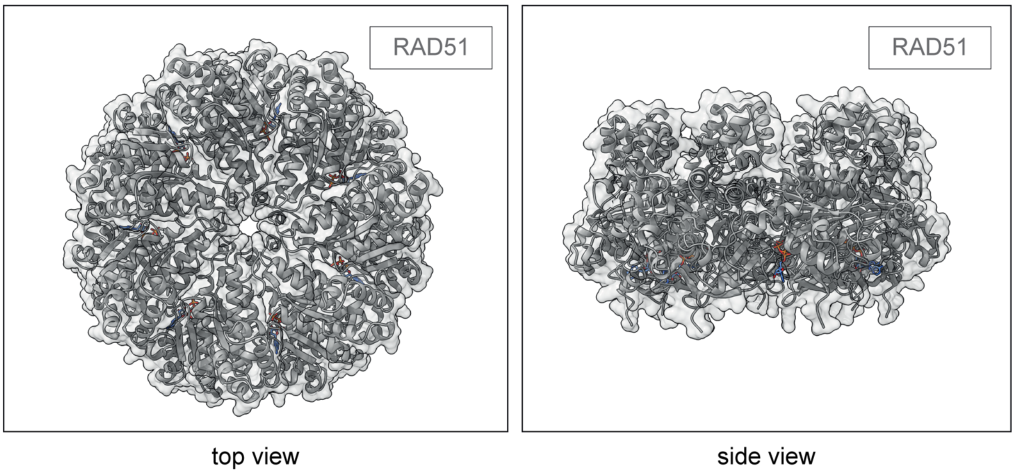


**B**

**RAD51(5)-CX3.ATP [AF3 model]**

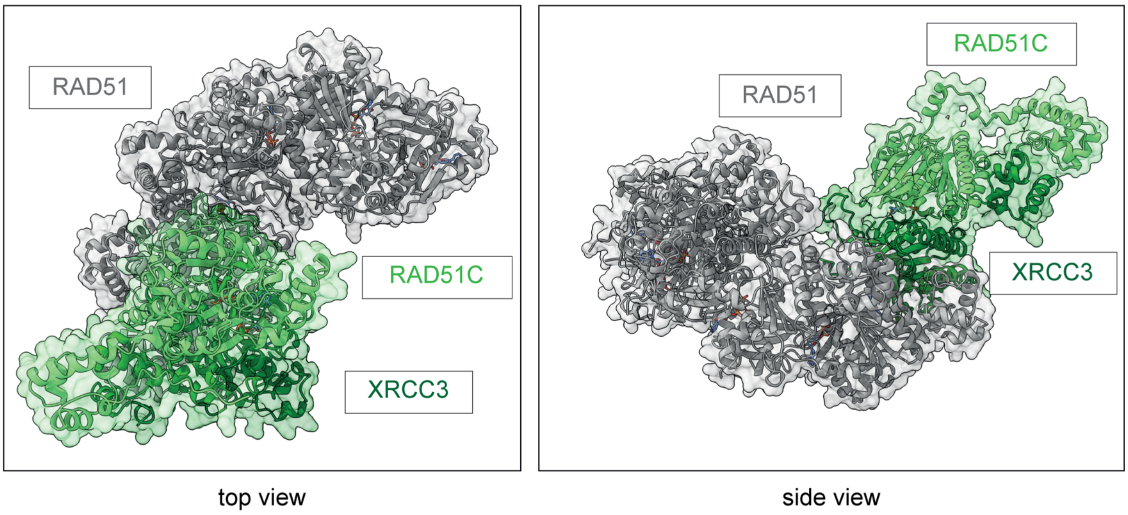


**Fig. S9: Comparison of Alphafold3 generated predictions of RAD51 oligomers or RAD51-CX3 complexes in the absence of DNA. (A)** AF3 prediction of ATP-bound RAD51 heptamer. **(B**) AF3 prediction of ATP-bound RAD51-CX3 heptameric complex.

***Supplementary methods:***

**Kinetic Data Analysis and Statistics.**

The kinetic data fitted globally with the KinTek Explorer program (KinTek, USA). This dynamic kinetic simulation program allowed multiple data sets to be fit simultaneously to a single kinetic model (Fig. S8C). Data fitting used numerical integration of rate equations from an input model searching a set of parameters applying the Bulirsch–Stoer algorithm with an adaptive step size that produces a minimum $\chi^2$ value calculated by using nonlinear regression based on the Levenberg-Marquardt method[63]. Residuals were normalized by sigma value for each data point. The standard error of the fitted parameters was calculated from the covariance matrix during nonlinear regression. The observable signal (*FI*) was defined as the sum of the contributions of all involved DNA species to the total Cy3 fluorescence. The Equation 1 was used to define the signal collected upon mixing CX3 with RAD51, where *f* scales the signal to concentration of CX3 and sensitivity of the measurement, factor *a* and *b* defines the relative change in fluorescence in forming RAD51.CX3$^0$ and RAD51.CX3 complexes, respectively. The Equation 2 was used to define the signal collected upon mixing RAD51 with DNA, where *f* scales the signal to concentration of DNA and sensitivity of the measurement, factor *a* defines the relative change in fluorescence upon RAD51 binding and forming DNA.RAD51$_n$ bound complexes, factors *b* and *c* define the relative change in fluorescence in forming filament DNA.*f*(RAD51$_n$) and its conformational rearrangement to DNA.*f*(RAD51$_n$)$^*$, respectively. The Equation 3 was used to define the signal of combined experiments mixing DNA with both CX3 and RAD51.

$$FI_{CX3.RAD51} = f \times (CX3 + a \times RAD51.CX3^0 + b \times RAD51.CX3)$$

*(Eq. 1)*

$$FI_{RAD51} = f \times (DNA + a \times (DNA.RAD51 + DNA.RAD51_n) + b \times DNA.f(RAD51_n) + c \times DNA.f(RAD51_n)^*)$$

*(Eq. 2)*

$$FI_{CX3,RAD51.DNA} = f \times (DNA + a \times (DNA.RAD51 + DNA.RAD51_n) + b \times DNA.f(RAD51_n) + c \times DNA.f(RAD51_n)^* + d \times DNA.Rad51.CX3 + e \times DNA.Rad51.CX3_n + f \times DNA.f(RAD51.CX3_n) + g \times DNA.f(RAD51.CX3_n)^*)$$

*(Eq. 3)*
